# Single-use, Metabolite Absorbing, Resonant Transducer (SMART) culture vessels for label-free, continuous cell culture progression monitoring

**DOI:** 10.1101/2024.01.27.577601

**Authors:** Yee Jher Chan, Dhananjay Dileep, Samuel M. Rothstein, Eric W. Cochran, Nigel F. Reuel

## Abstract

Secreted metabolites are an important class of bio-process analytical technology (PAT) targets that can correlate to cell condition. However, current strategies for measuring metabolites are limited to discrete measurements, resulting in limited understanding and ability for feedback control strategies. Herein, we demonstrated a continuous metabolite monitoring strategy using a single-use metabolite absorbing resonant transducer (SMART) to correlate with cell growth. Polyacrylate was shown to absorb secreted metabolites from living cells containing hydroxyl and alkenyl groups such as terpenoids, that act as a plasticizer. Upon softening, the polyacrylate irreversibly conformed into engineered voids above a resonant sensor, changing the local permittivity which is interrogated, contact-free, with a vector network analyzer. Compared to sensing using the intrinsic permittivity of cells, the SMART approach yields a 20-fold improvement in sensitivity. Tracking growth of many cell types such as Chinese hamster ovary, HEK293, K562, HeLa, and *E. coli* cells as well as perturbations in cell proliferation during drug screening assays were demonstrated. The sensor was benchmarked to show continuous measurement over six days, ability to track different growth conditions, selectivity to transducing active cell growth metabolites against other components found in the media, and feasibility to scale out for high throughput campaigns.

## Introduction

Process analytical technologies (PAT) for continuous monitoring of cell cultures enable optimization and feedback control in biomanufacturing, as well as provide insight to understand process effects on cells.^[1–5]^ Such tools are useful in traditional biomanufacturing to track progress of cell factories used to generate end use products (e.g., antibodies, small molecules); however, with the advances in therapeutic cells as “living medicines,” there is an even greater need for PATs to control and understand the health of cells in these complex bioproduction systems. For instance, manufacturing of autologous CAR T cells can have highly variable expansion rates and phenotypes due to different genetic and treatment histories of patients.^[6–9]^ Current process parameters continuously monitored in cell manufacturing are limited to physical parameters such as temperature, pH, and dissolved oxygen concentration within the culture media.^[10–13]^ Although these parameters can be used to maintain a healthy growth environment, these measurements do not accurately indicate the growth progression of cells. Monitoring cell proliferation directly would inform feeding, stimulation, and harvest decision time points.

Currently, most cell expansion processes in manufacturing settings are being monitored by intermittent sampling and analyzed in an external device (e.g. cell counter, microscope). In discovery settings, fluorescence approaches can be used to provide a high signal to background ratio on labeled cell parts (e.g. nuclei) and the images can be automatically segmented for high-throughput measurements of cellular properties. Although microscopic techniques can provide real-time information over a few hours, they are not readily applicable due to complex instrumentation requirement and difficulty in coupling with industrial reactors.

For continuous measurement of cell growth, electric field based methods can be used to assess cell growth; these are non-destructive and can be integrated into traditional vessels. For instance, bio-capacitance probes, such as those offered by Aber and Hamilton, can be inserted into the sampling ports of large-scale stirred bioreactors (>250 mL).^[14–18]^ These larger probes are not well-suited for use in small reactors. For some smaller bioreactors, electric cell-substrate impedance (ECIS) technology, such as the real time cell analyzer (xCELLigence) can be used to measure adhesion, transendothelial electrical resistance, and growth progression of adherent monolayers.^[19–26]^ Another electric field-based approach is inductive-capacitive (LC) sensors, also known as resonant sensors. Like bio-capacitance probes, resonant sensors are affected by changes in local permittivity and can be used to monitor presence of cells while neglecting effects from cell debris. Studies have been performed to investigate their feasibility to detect live cells, but these include microfluidic sampling channels to increase sensitivity.^[27–31]^ To date, tracking cell growth directly in traditional culture vessels with resonant sensors has not been a reliable approach.

While optical and electrical methods provide a non-invasive way of probing biological systems, measuring secreted metabolites represents an alternative approach that can have minimal impact on cells while providing implications to many biological processes.^[32–36]^ For instance, lactate and glutamate concentrations are often measured from the sampled culture media due to their correlation with metabolism, and thereby cell growth. In addition, colorimetric assays involving tetrazolium salts are used to reflect metabolic activity via reduction reactions by mitochondrial enzymes. However, metabolite measurements often involve an enzymatic reaction that makes continuous monitoring challenging. Other analytical tools such as mass spectrometry, Raman^[37–39]^ and Nuclear Magnetic Resonance spectroscopy^[40–43]^ have recently been investigated and developed for metabolites and other biomolecules analysis. However, these methods typically require a bypass line from bioreactors and depend strongly on calibration curves, posing challenges for integration into diverse culture systems.

Another strategy for metabolite sensing is coupling a biological or synthetic receptor to a sensitive transducer. Biological receptors such as antibodies and aptamers can be engineered to have a strong binding affinity toward target metabolites. However, their long-term stability within the culture environment are limited by enzymatic degradation and they are also not readily sterilizable by conventional techniques. Synthetic receptors, such as cross-linked polymers, have superior stability and have demonstrated affinities to organic molecules, finding use in passive, environmental air sampling^[44]^. In some cases, these absorbed molecules can plasticize the polymer, changing its physical properties, such as with conducting polymers and gas sensing.^[45,46]^ Molecular imprinted polymers is an approach to engineer higher affinities to target molecules in polymer receptors.^[47]^ Polymers as synthetic receptors have not been widely applied in bioproduction, likely due to insufficient selectivity, especially in complex culture media. Therefore, targeting a class of molecules rather than a single metabolite target, might be a more fruitful approach, especially when tracking a global parameter, such as average cell growth.

In this work, we present the development and applications of a resonant sensor coupled to a signal enhancing polyacrylate to transduce cell growth through extracellular metabolites. First, we evaluated the sensitivity of resonant sensors to cells without the polymer, where the permittivity contrast between live cells and surrounding media is measured directly resulting in a smaller sensitivity. Next, we explored the utility of a single-use, metabolite absorbing, transducing (SMART) material to increase the sensitivity. We employed a polyacrylate adhesive film as the SMART material and investigated its physical property responses when screened with primary and secondary metabolites. We demonstrated the SMART sensor capability to continuously monitor cell growth and death with a sampling rate of 10 minutes for up to 5 days where a steady-state signal was observed. We show how fit parameters to the dynamic responses can be used to correlate different culture conditions including growth variations caused by various start cell concentrations as well as drug screening responses. Through this innovation, a continuous cell progression monitoring platform that is compatible with most single-use bioreactors is shown, showing wide application in biomanufacturing and biological studies.

## Results and Discussion

### Resonant sensors directly reporting cell growth – operating principles

Resonant sensors, also known as passive LC sensors, are wirelessly interrogated through inductive coupling. The system is comprised of a patterned sensor coil^[48]^ and a reader antenna connected to a reader (vector network analyzer or custom impedance board)^[49]^ that sweeps through frequencies and observes changes in reflected or transmitted power. The sensor coil has an intrinsic inductance and parasitic capacitance (LC), which upon inductive coupling, oscillates at a specific resonant frequency. These sensors are tuned to operate in the short-wave radio frequency spectrum to balance sensor size and capability to transmit longer distances through cell growth media. When sweeping and measuring the impedance through the reader coil, the resonant frequency is indicated by a loss in reflected power^[50]^ measured in s-parameters, e.g., S_11_ (Figure 1b). Variation in permittivity proximal to the sensor in response to the target analyte is a common sensing modality, as this has a large effect on resonant frequency.^[51]^

**Figure 1.**
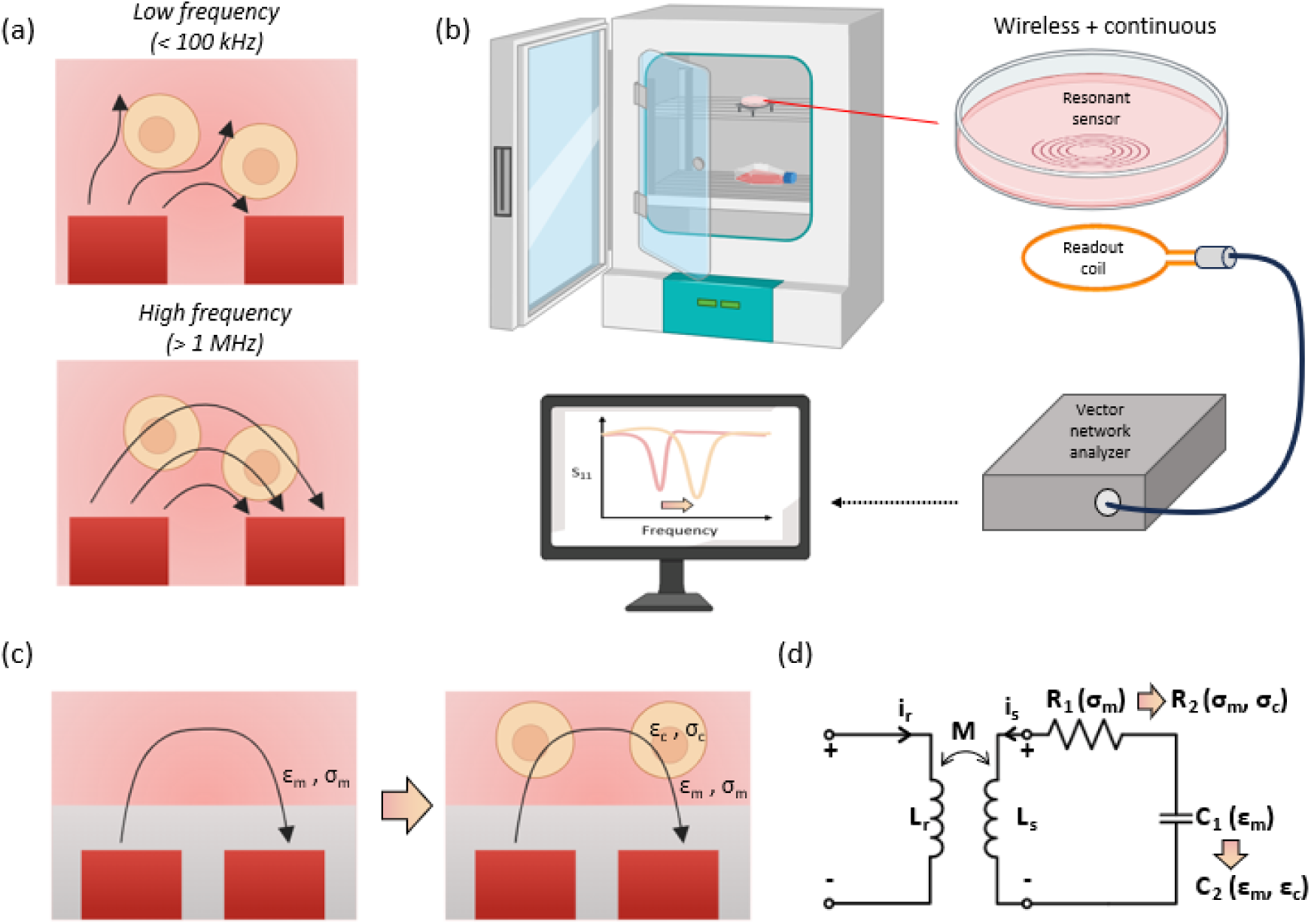
Sensing principle of a resonant sensor coupled directly to cell growth. (a) Schematic of electric field behavior on cells at different frequencies showing that electric field does not penetrate through cells at low frequencies and only does so at higher frequency where β-dispersion occurs. (b) Schematic of the resonant sensor platform for *in vitro* cell growth monitoring. The resonant sensor sits on the bottom of a culture dish while a reader coil inductively couples with the resonant sensor outside of the dish. Illustration of reflectance (*S*_*11*_) response when swept over a range of frequencies during the cell growth, depicting an increase in frequency of the reflectance minimum caused by decrease in relative permittivity local to the cell. (c) Schematic of operating principles of resonant sensor in biological cultures. *ε*_*m*_ and *ε*_*c*_ represent the permittivity of culture media and cell, respectively, whereas *σ*_*m*_ and *σ*_*c*_ represent the conductivity of culture media and cell, respectively. Live cells around the sensor induce a difference in permittivity around the sensor which can indicate growth. (d) Equivalent circuit of resonant sensor sensing system. *L*_*r*_ and *L*_*s*_ are the inductance of reader coil and resonant sensor, respectively. Inductive coupling between the two coils results in the mutual inductance, *M*. *C*_*1*_ and *C*_*2*_ represent the capacitance change resulting from permittivity variation in the absence and presence of cells, respectively. *R*_*1*_ and *R*_*2*_ represent the resistance change resulting from conductivity variation in the absence and presence of cells, respectively.

Two modes of signal transduction are possible when applying resonant sensors to cell culture sensing (Figure 1a). First, cell membranes are composed of a low conductivity phospholipid bilayer that separates the intercellular contents from the surrounding medium, functioning as a capacitor at low frequencies (< 100 kHz). Since the electric field is impeded by the cells, measuring the impedance between two electrodes can reflect the effects introduced by cells, as in the case of ECIS. Similarly, the presence of cells can alter the electromagnetic fields around the resonant sensor and change the resonant frequency. However, as lower resonant frequencies generally require a larger inductive coil, the sensing area also proportionally increases. This results in a negligible effect of the capacitance introduced by the cells when compared to the bulk environment. Therefore, the cell as a capacitor principle is not a good strategy for a passive LC sensor. However, as the frequency increases (> 100 MHz), the cell membrane no longer functions as an insulator due to β-dispersion. At this frequency range, cell conductivity decreases relative to the bulk and the current begins to penetrate through the cytoplasm, resulting in a resonant frequency that is sensitive to the intercellular permittivity, *ε_c_*. Fortunately, the intercellular permittivity differs from the culture media and is dependent on cellular properties such as water content and cell size. Therefore, the cell growth can be detected by starting with media permittivity and conductivity (*C_1_*(*ε_m_*), *R_1_*(*σ_m_*)) and progressing to that of a mixture of cells and media (*C_2_*(*ε_m_*,*ε_c_*), *R_2_*(*σ_m_*,*σ_c_*)), shifting the resonant frequency throughout the cell culture progression (Figure 1c,d).

### Thin coated resonant sensor exhibits lower sensitivity towards cell growth

Resonant sensors are highly sensitive to the surrounding conductivity. A high conductivity environment can reduce the quality factor of the sensor, as in the case of culture media. Therefore, an insulating layer is needed to preserve the sensor signal. Using simulation (Ansys HFSS), the sensor system was investigated to generate design guidelines to achieve maximum sensitivity towards the cells. A thin (3 μm) layer above the sensor and the insulator was used to simulate a confluent layer of cells (Figure S1). The resulting resonant frequency from different permittivity of the thin cell layer was simulated (Figure 2a). The change in resonant frequency increases when the permittivity of the cell layer decreases due to the stronger permittivity contrast from surrounding media. In addition, a thicker insulator (sensor coating) reduces the sensitivity of the sensor, as noted by the steeper curve at lower insulator thickness. A coating thickness of 50 μm halved the sensor gain of a 10 μm coating. This simulation shows there is a balance between increased insulation thickness needed to preserve sensor signal strength and keeping it thin enough to detect the cells and maintain a large sensor gain.

**Figure 2.**
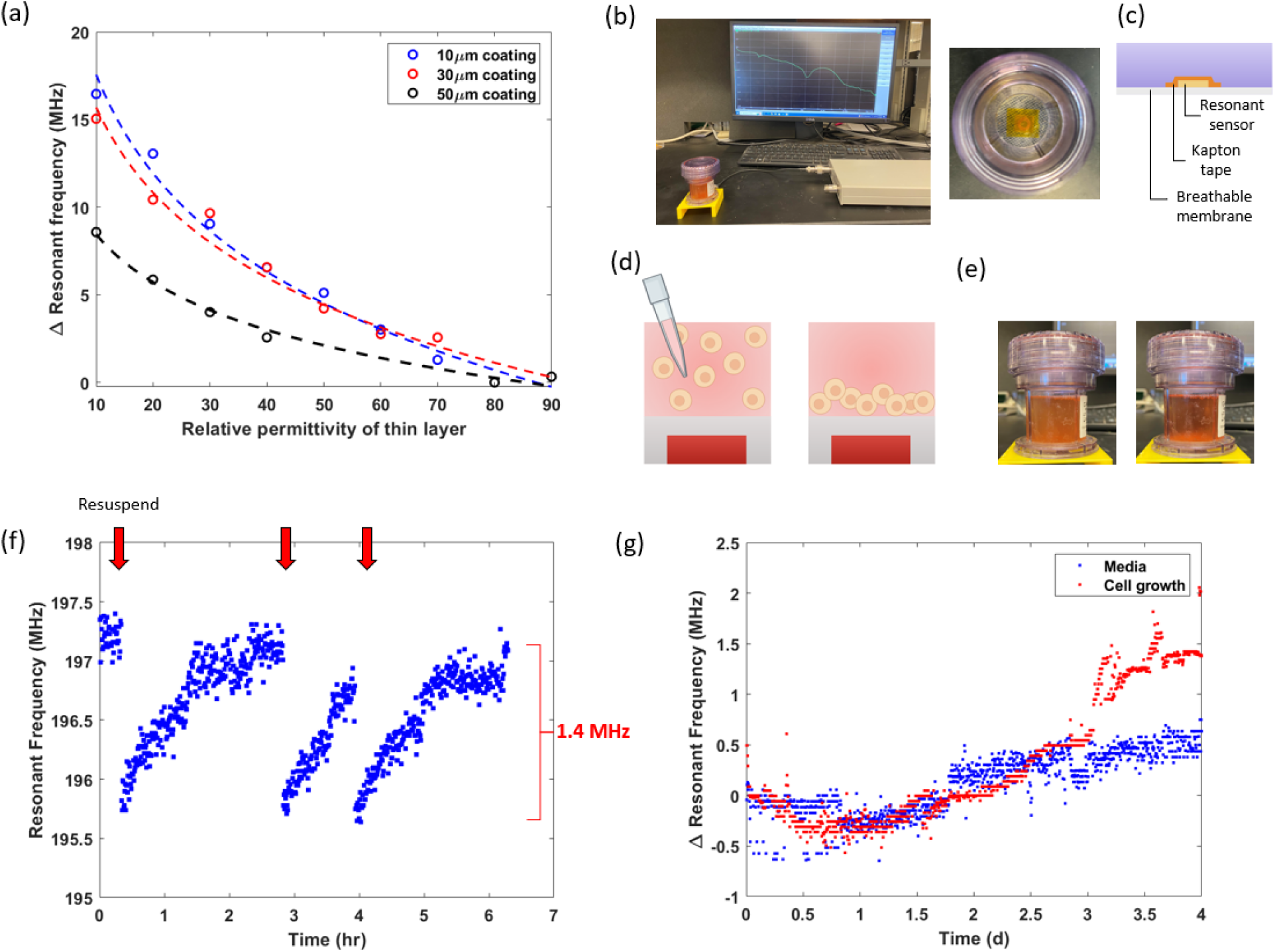
Investigation of resonant sensor sensitivity to live cells caused by intrinsic cell permittivity contrast from the surrounding media. (a) Simulation of resonant sensor responses to permittivity variations of a thin 3 μm film around the sensor at different insulator coating thicknesses. An insulator protects the resonant sensor from shorting the signal. (b) (Left) Image of the experiment setup for cell resuspension experiment. (Right) Top view of the G-Rex10 culture flask containing resonant sensor. (c) Schematic illustration of the cross-sectional side view of G-Rex10 to show the layered structure. (d) Schematic illustration of the experimental design for rapid investigation of resonant sensor sensitivity towards live cells. Gradual resuspension process of confluent K562 cell culture was used to simulate cell growth. (e) Image of G-Rex10 containing dispersed cells (left, cloudy) and settled cells (right, clear). (f) Resonant frequency response to three resuspension events at insulator thickness of 63.5 μm. Red arrows indicate each resuspension event followed by settling of the cells. (g) Resonant frequency response without cells (control) and with K562 cells over 4 days of culture using G-Rex membranes as insulator (thickness = 200 μm), showing limited signal above control.

The resonant sensor form factor and design can affect the sensor sensitivity by changing its sensing region. For instance, resonant sensors can be designed to have different overall diameters with similar resonant frequencies (keeping the inductive coil as the same overall trace length); however, a smaller sensor diameter generates a smaller electric field and is more sensitive to the permittivity within a smaller region. This feature is desired when the target-induced permittivity change is confined locally around the sensor as the cells settle in the electric field generated by the sensor. With the design rules described, we have selected a resonant sensor with 1 cm diameter. It was noted that further reducing the form factor reduced the interrogation step-off distance (antenna to sensor) and the coupling efficiency (increased signal to noise).

As a proof of concept, the sensor sensitivity to the changing environment permittivity at micron scale under different sensor coating (insulation) thicknesses was studied. As a proxy to live cells, we suspended polyethylene beads (25 μm diameter, *ε_polyethylene_* ≈ 3) into a buffered solution and resuspended the mixture, simulating cell growth as the beads gradually deposited into a 300 μm multilayer. The sensor achieved up to 7 MHz shift at an insulation thickness of 100 μm and below, whereas the response of a 200 μm coated sensor reduced to 2 MHz (Figure S2, S3). While the reduced sensitivity at thicker insulation thickness is expected due to weaker permittivity effects, an unexpected finding occurred by further reducing the insulation thickness below 100 μm and also observing reduced reflected power (*S_11_*) and smaller resonant frequency shift. This is mainly caused by decreased resonant frequency at lower insulation thickness (limiting the extent of shift) and depicts an optimal sensor coating thickness when measuring cells directly on the resonant sensor.

To validate and identify the sensitivity towards actual cell culture, we performed a similar resuspension experiment with a fully expanded K562 cell culture in a breathable static cell culture vessel (G-Rex10, Wilson Wolf) (Figure 2b,c,d,e). The tall static culture vessel was used to provide sufficient liquid height to minimize the signal effect of cells when dispersed. Similar to the beads and modeled experiments, we observed a sudden decrease in the resonant frequency (1.4 MHz) immediately after resuspending the fully grown cells (Figure 2f). This is consistent with the reported cell relative permittivity of 30-70^[52,53]^, as the culture media replacing the cells has a higher permittivity (*ε_media_* ≈ 80), thereby increasing the capacitance and reducing the resonant frequency immediately. The resonant frequency gradually returns to the initial frequency as the cells slowly redeposit after about 90 minutes of settling (Fig 2f). A lower sensitivity compared to the beads was expected due to different diameter and permittivity (*ε_polyethylene_* ≈ 3).

After confirming sensitivity towards the cells, we proceeded with monitoring the cell growth in its native *in vitro* culture environment. We cultured the cells in the G-Rex10 and measured the resonant frequency change over 4 days after cell inoculation (Figure 2g). Unfortunately, the resonant frequency response was 0.9 MHz from noise level of the control vessel (no cells inoculated) due to the exposed sensor being susceptible to many background factors such as humidity and temperature changes in the incubator, resulting in a background signal drift that covered the majority of the sensor response especially at low cell concentrations. The sensor stability could be further improved with a back coating that shields from environmental fluctuations, although this would limit the signal strength and the response would still be limited to a few MHz shift. We therefore put our attention on advanced materials to increase the sensor gain (extent of frequency shift to equivalent cell growth).

### Study of acrylic adhesives for mechanical sensitivity to secretomes

As shown above, cell growth can be monitored directly with resonant sensors using the intrinsic permittivity difference of cells to media, however this has limited sensitivity due to small changes in cell internal permittivity and the requirement to keep sensors proximal to the cell growth area. After initial testing we utilized a polyacrylate adhesive film as the biomolecule responsive material to amplify the signal due to its solvent resistance (will not lift off during long cell growth experiments) and extent of published plasticizers. Polyacrylate has a tunable modulus when mixed with plasticizers such as terpene and certain hydrocarbon resins.^[54]^ Since biological cells are known to secrete a wide range of hydrocarbon molecules, we employed an agnostic screening approach (targeting a range of molecules with a certain functional group, rather than a single molecule) by examining the polymer property in response to biomolecules found in cell culture. The impregnated polymer likely has a small change in relative permittivity, but we realized that the resulting mechanical change could be transduced into a larger resonant frequency shift; the polyacrylate adhesive film was applied onto a wire wound resonant sensor in a dish, intentionally creating voids around the wire (Figure 3b). As the polyacrylate film absorbs biomarkers, it plasticizes and softens, closing out the air voids, which creates a strong permittivity contrast (from *ε_air_* ≈ 1 to *ε_adhesive_* ≈ 5 and *ε_media_* ≈ 80) when the mechanical property changes (Figure 3a, right). This resulting prototype is abbreviated as a single-use metabolite absorbing resonant transducer (SMART) system.

**Figure 3.**
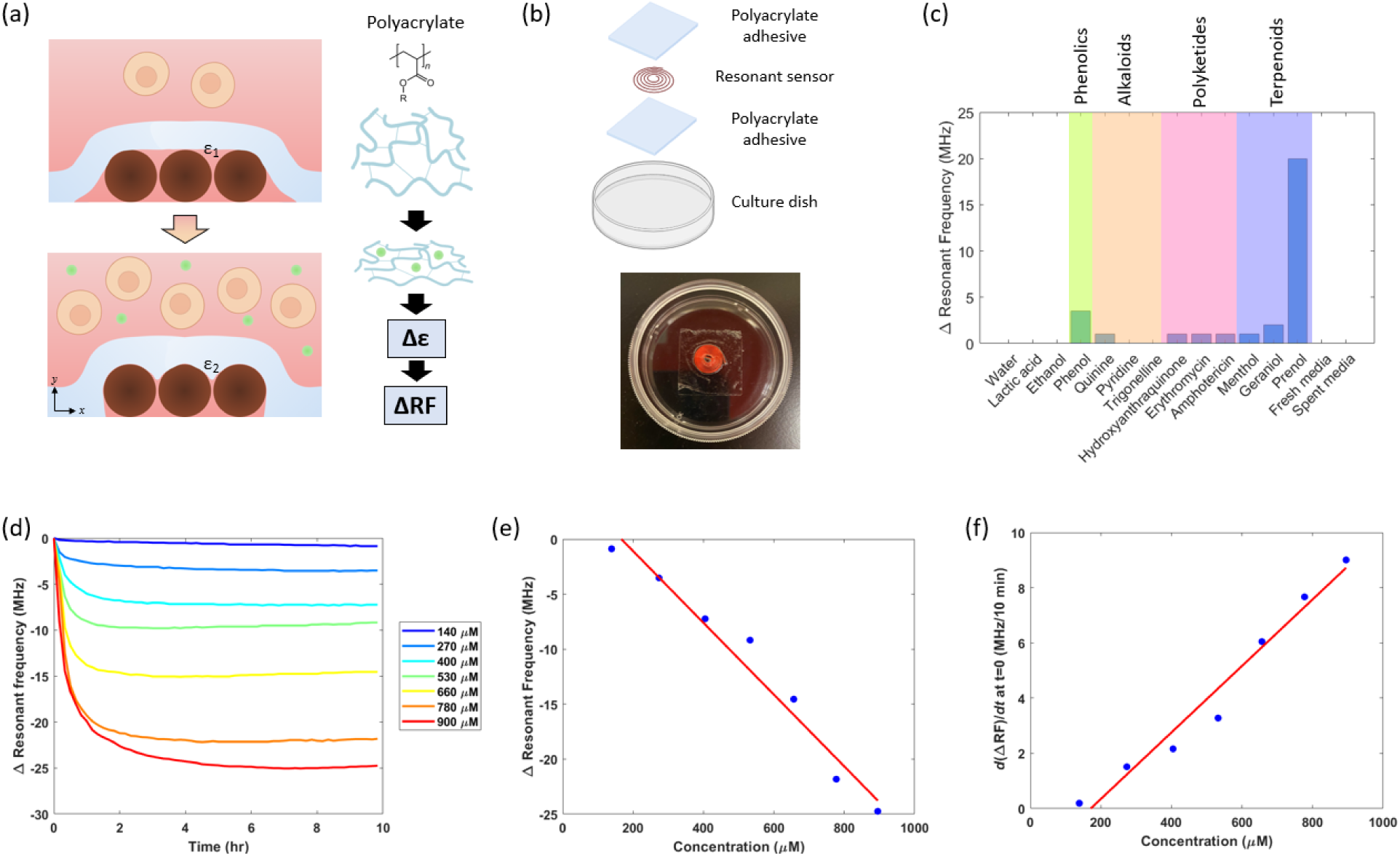
Working principle and selectivity screening of SMART dish. (a) Schematic illustration of the working principle of polyacrylate film: softening upon exposure to growth metabolites inducing a permittivity contrast detected by the resonant sensor. (b) Exploded view and image of the SMART dish. (c) Resonant frequency response to different classes of metabolites including the fresh and spent media that includes mixture of all primary and secondary metabolites. (d) Temporal resonant frequency response to different concentrations of prenol. (e) Total resonant frequency change at steady state at different prenol concentrations. Data was fitted with a linear model as shown by red line. (f) Maximum rate of resonant frequency response at various prenol concentrations. Data was fitted with a linear model as shown by red line.

To examine the sensitivity and selectivity of the SMART setup towards different classes of biomolecules, we evaluated the signal response resulting from biomolecules containing different functional groups. An ideal sensor will be non-responsive to proteins and most primary metabolites that are abundantly present in fresh media as they would interfere with cell growth monitoring by creating an irreversible mechanical response upon initial exposure to these common molecules present in media. Secondary metabolites, on the other hand, are produced in proportion to the biological processes within the cells that can be used to indicate cell growth. We selected several molecules of different molecular weights and solubility from each secondary metabolite class (phenolics, alkaloids, polyketides, and terpenoids) for screening (Figure 3c). Phenolics and terpenoids, specifically phenol and prenol, resulted in a higher sensitivity compared to other classes, giving a 3.5 and 20 MHz signal change, respectively. Notably, the molecules that exhibited a higher sensitivity contain alkenyl and hydroxyl functional groups. Geraniol (another terpenoid molecule) resulted in a 2 MHz response; however, the signal continuously changed after 10 hours while the signals from other molecules typically plateaued, suggesting a diffusion-limited response due to a longer carbon chain and lower solubility (Figure S4). Menthol, which is absent of unsaturated carbon bonds, exhibited only 1 MHz response.

To examine the effect from other free flowing biomolecules, we performed further screening using commonly known secreted biomolecules such as nucleotides, proteins, and extracellular vesicles. The SMART dish was first incubated with culture media for about 20 hours to allow signal equilibration resulting from the media diffusion into the adhesive. The initial addition of media to SMART resulted in a signal change, but eventually reaches steady-state (Figure S5). Further addition or replacement of the culture media did not result in an appreciable signal response (<2 MHz), suggesting minimal effect from amino acids and ions that are abundantly present. Similarly, addition of serum did not cause a signal change, suggesting that the SMART is not selective to proteins and nucleic acids (Figure S6). For sensitivity towards extracellular vesicles or lipids, liposomes were added and resulted in no signal change (Figure S7). These implied that the sensor signal will not be affected by media components and most undesired molecules. Importantly, we observed that the spent media (end of cell growth) did not contain enough metabolite to cause a shift, indicating that there would need to be a transient buildup of the metabolite to induce a change – again a desirable feature as we want the sensor to respond to active cell growth or cessation of growth.

Noticing exceptional sensitivity towards molecules containing alkenyl and hydroxyl groups such as prenol, we performed further investigation under various prenol concentrations ranging from 140 to 900 μM. Although the selected concentrations are higher than typical metabolites concentration found in culture media, we hypothesized that this concentration could be achieved cumulatively throughout the long-term growth of cell culture. We observed increasing resonant frequency shift with increasing prenol concentration, reaching steady state after several hours (Figure 3d). When extracting the end point resonant frequency shift at 10 hr, we observed a linear relationship with the initial prenol concentrations (Figure 3e). In addition, the rate of frequency shift at maximum (time ≈ 0) also increases linearly at increasing prenol concentrations (Figure 3f), suggesting that the concentration of absorbed prenol and the rate of absorption varies depending on the prenol concentration around the polyacrylate film. The diffusion reaction of the system can therefore be simply modeled as follow:

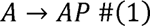

Where A is the prenol molecules in the bulk solution and AP is the embedded molecules within the polymer film. Assuming the concentration of AP is reflected by the resonant frequency, the rate of formation of AP can be described as

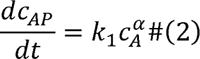

where k_1_ is the rate constant. According to linear correlation shown in Figure 3f, we can infer that *α*=1. Applying mass balance (*c_A_* = *c_A0_* - *c_AP_*) and solving the linear equation gives

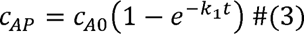

Letting the resonant frequency to be correlated with c_AP_ by a constant m, the equation can be rewritten as:

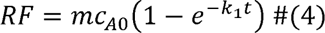

Evaluating equation 4 as t➔ ∞ (steady state), the constant m can be determined from the slope obtained in Figure 3e, m = -0.0326 MHz/μM. Subsequently, k_1_ can be determined by taking the derivative of equation 4 and compared to the slope in Figure 3f, k = -0.037 min^-1^. Note that the rate constant is negative due to the decreasing resonant frequency. This model matches with the exponential decay response curve obtained from prenol plasticizing into polyacrylate (Figure 3d) and suggests concentration-dependent absorption. This equilibrium behavior is critical for the continuous measurement.

### Validation of polymer mechanical responses to biomolecules

We next validated that the exposure of this film to secreted biomolecules caused a mechanical change, as we hypothesized in our mechanism (Fig 3a). We layered the adhesive film on top of a polyethylene terephthalate film with patterned channels to create air voids (Figure 4a). As the adhesive interacts with the secreted biomolecules in the presence of cells, changes in the mechanical property can cause the adhesive film to sink into the voids (Figure 4b). In the setup where cells were introduced, we indeed observed that the polymer film conformed into the air voids whereas the film that was only incubated in the culture media did not (Figure 4c,d). This confirmed the mechanical response to cell culture and indicated a permittivity variation resulted from this response.

**Figure 4.**
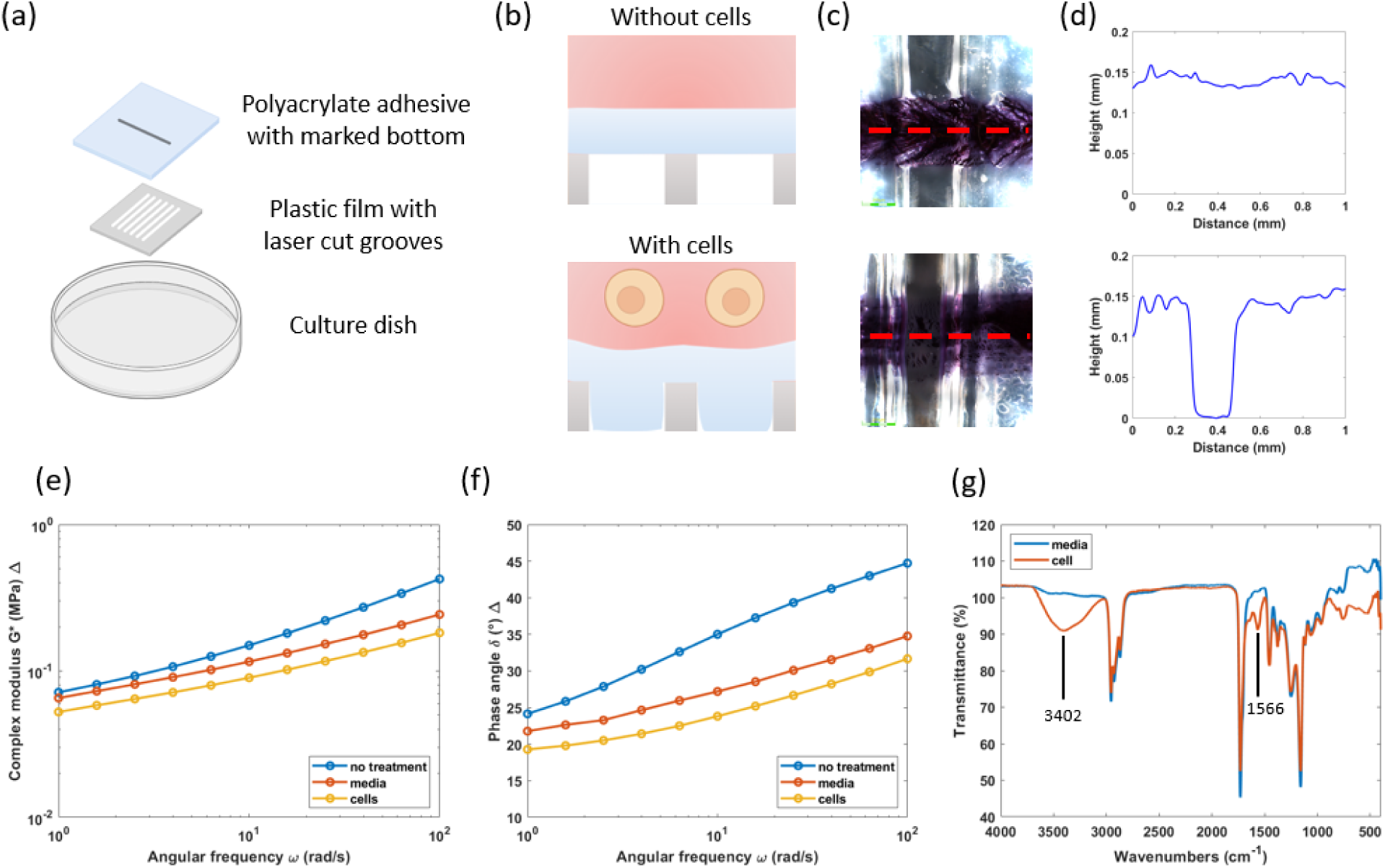
Validation of mechanical response and characterization of polyacrylate film to metabolite exposure. (a) Exploded view of the experimental setup used to validate the mechanical response of polyacrylate film. Black trace across the PET cut well was marked under the polyacrylate film (above PET film) to provide contrast for height profiling. (b) Schematic illustration of side cross-sectional view of the conformation of polyacrylate film into the voids in response to cell growth. (c) Top-down microscopic view of the setup in (a) after 4 days of incubation without cells (top) and with cells (bottom). Red dotted line represents the cross section at which the height profile was obtained with a digital metrology microscope. (d) Height profile of polyacrylate adhesive without cells (top) and with cells (bottom). (e,f) Rheological characteristics of polyacrylate without any treatment, treatment with media, and treatment with cells. (g) FTIR spectra of polyacrylate treated with media and with cells.

To further evaluate the mechanical properties, rheological information was obtained for polyacrylate with and without exposure to cells. After exposure to fresh media or cell suspension over 4 days, polyacrylate in both conditions resulted in a lower complex modulus (*G\**) and phase angle (*δ*) compared to the untreated polyacrylate (Figure 4e,f). Treated polyacrylate in cell suspension exhibited a lower *G\** and *δ* compared to media only exposure, suggesting that the stiffness decreased and a more elastic behavior in the cell-exposed polyacrylate due to a higher degree of plasticization from the secretion. Differential scanning calorimetry and thermogravimetric analysis of polyacrylate were also performed to examine the thermal properties but exhibited similar characteristics in both conditions (Figure S8).

We then performed molecular characterization on polyacrylate to identify the molecules and chemical properties using Fourier transform infrared spectroscopy (FTIR) and liquid chromatography-mass spectrometry (LCMS). It is notable that the polyacrylate adhesive turned slightly yellowish upon treatment with cell growth. For the FTIR spectra, the cell-treated polyacrylate resulted in two additional characteristic peaks compared to the media-treated at 3402 and 1566 cm^-1^ (Figure 4g). The intense broad peak at 3401 cm^-1^ is indicative of multiple hydrogen-bonded interactions due to the presence of a free hydroxyl group, which is not uncommon in the terpenoid class. Further, a weak but broad peak at 1566 cm^-1^ can arise due to C(sp2)-C(sp2) stretching modes. Typically, higher frequencies (1600-1650 cm^-1^) are more common for alkenes; however, extended conjugation (often found in carotenoids) can often reduce the energetics of the stretch, causing a bathochromic shift. This is consistent with the greater response observed with the terpenoids class (Figure 3c), demonstrating a higher affinity toward molecules containing these functional groups, though not all the molecules may specifically be terpenoids. To validate this, the polyacrylate was swollen with acetone to extract the embedded molecules and analyzed using LCMS. Both the media-treated and cell-treated polyacrylate resulted in many features. Among the differentially expressed molecules in the cell-treated polyacrylate, most of the molecules contained hydroxyl and alkenyl/benzyl functional groups (See supporting materials for a complete least of upregulated molecules).

### Polyacrylate adhesive as a signal amplifier for cell growth when coupled to resonant sensors

We next investigated if the SMART system could monitor cell culture progression using the mechanical change as a transducing parameter to convert the cell growth metabolites into an electrical signal through the resonant sensor. HeLa cells were selected as the model as the adherent property could reflect cell physiology. A reader coil integrated with a frugal microscopic imaging system (Figure S9) was built to synchronously examine the sensor responses and cell growth images (Figure 5a,b). After 4 days of culture, we observed over 30 MHz of resonant frequency shift (Figure 5c), a sharp improvement to the 1.4 MHz response from sensing using intrinsic cell permittivity. The dynamic sensor response illustrated a sigmoidal curve, where a slight decrease in resonant frequency was first observed in the first 45 hours after cell seeding, followed by a dramatic decrease that lasted for about 30 hours, and finally plateaued. Time-lapse microscopic images of the cells confirmed the proliferation of cells throughout the experiment (Figure 5d). At 0 hr, HeLa cells were seeded and began attaching to the culture dish and the adhesive. Cell expansion was not prominent in the first 24 hr, maintaining at around 70% confluency, as shown in time stamp 1. At time stamp 2 (48 hr), the resonant frequency entered the exponential phase, and the cells expanded rapidly to form a fully confluent monolayer. At time stamp 3 (65 hr), the resonant frequency response began slowing while the cells formed a more compact monolayer. At time stamp 4 (94 hr), the resonant frequency plateaued, and the cells remained compact and slowly built a secondary layer. This strongly indicates that the resonant frequency change correlates to cell proliferation.

**Figure 5.**
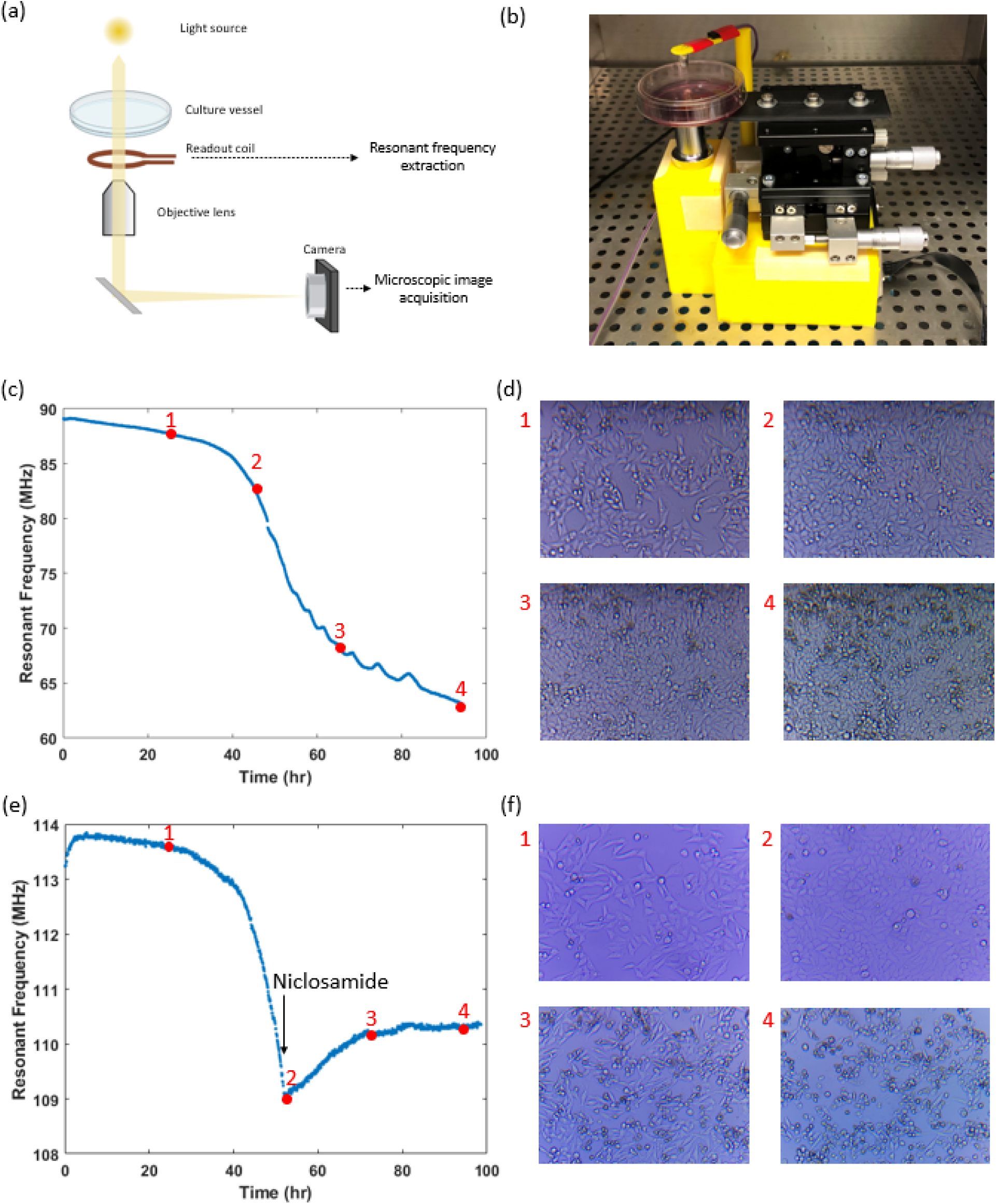
SMART response to cell growth and death. (a) Schematic illustration of the wireless readout system consisting of a readout coil for resonant frequency measurement and a frugal microscopic imaging system. (b) Image of the operating SMART system with ability for simultaneous resonant frequency and microscopic image acquisition. (c) Resonant frequency response over 4 days of HeLa cells culture. (d) Microscopic images of cell culture corresponding to different time stamps noted in (c). (e) Resonant frequency response with an interruption to the cell growth. Niclosamide was added at 52hr to interrupt cell growth. (f) Microscopic images of cell culture corresponding to different time stamps noted in (e).

The sigmoidal response is a result of metabolite accumulation in the polyacrylate and can be attributed to the typical growth curve of cells (increased metabolite secretion with more cells), albeit in reverse magnitude. As shown in Figure 3c, the spent media alone did not result in a similar resonant frequency shift. Media exchange during the cell growth also demonstrated a similar sigmoidal response (Figure S10). These results show that the corresponding metabolites can undergo degradation and the SMART responses reflect the cumulative measurement of metabolites, omitting needs for high detection limit.

### Effects of cell growth inhibitors on sensor response

Having confirmed signal response to cell growth, we next characterized response to cell death or non-proliferation. SMART dishes were cultured with HeLa cells and the resonant frequency and microscopic images were obtained as described above. After 52 hours, once the exponential growth phase was transduced with the rapid decrease in resonant frequency, niclosamide (an antihelminthic drug that interrupts the canonical Wnt signaling) was administered into the cell culture (30 μM final concentration). the resonant frequency stopped decreasing within a minute upon the drug administration (Fig 5e). Simultaneous microscopy further confirmed that the cells were proliferating, reaching a confluent state prior to introducing the drug and then stopped proliferating after drug addition (Fig 5f). Twenty hours after drug addition, the cells began to shrink and detach, indicating cell death had occurred. Another twenty hours later, most of the cells have detached from the dish. These results indicate that the transducing metabolites stopped being synthesized when cell expansion was disturbed and that was sufficient to turn off the SMART response. This indicates that the sensor can dynamically track both progression and cessation of proliferation. In addition, while the microscopic images took time to show gradual shrinkage of the cells, the SMART dish was able to inform perturbation instantaneously. The gradual increase in resonant frequency after the drug addition is still being studied, and could be due to a small amount of reversibility in the film stiffness (leaching out some of the bound metabolites). Generally, the SMART dish generates an irreversible response that is complementary to single-use reactors.

### High throughput SMART readout system for tracking growth rates and correlating with cell confluency by microscopy

In-line sensors can provide real-time information of biological culture, but should be scalable for high-throughput measurement. The resonant sensor system holds an advantage in this aspect as a multiplexer can easily be integrated to the VNA to increase the measurement throughput without the need of complex system (Figure 6a,b). For instance, high-throughput microscopic imaging system requires a highly precise three-dimensional translation stage to acquire images. This is not required in the SMART prototype as the sensor measures the bulk culture property, driving the capital cost down in the SMART system. Additionally, the SMART dish demonstrated no significant effect on the cell growth and viability (Figure 6c,d).

**Figure 6.**
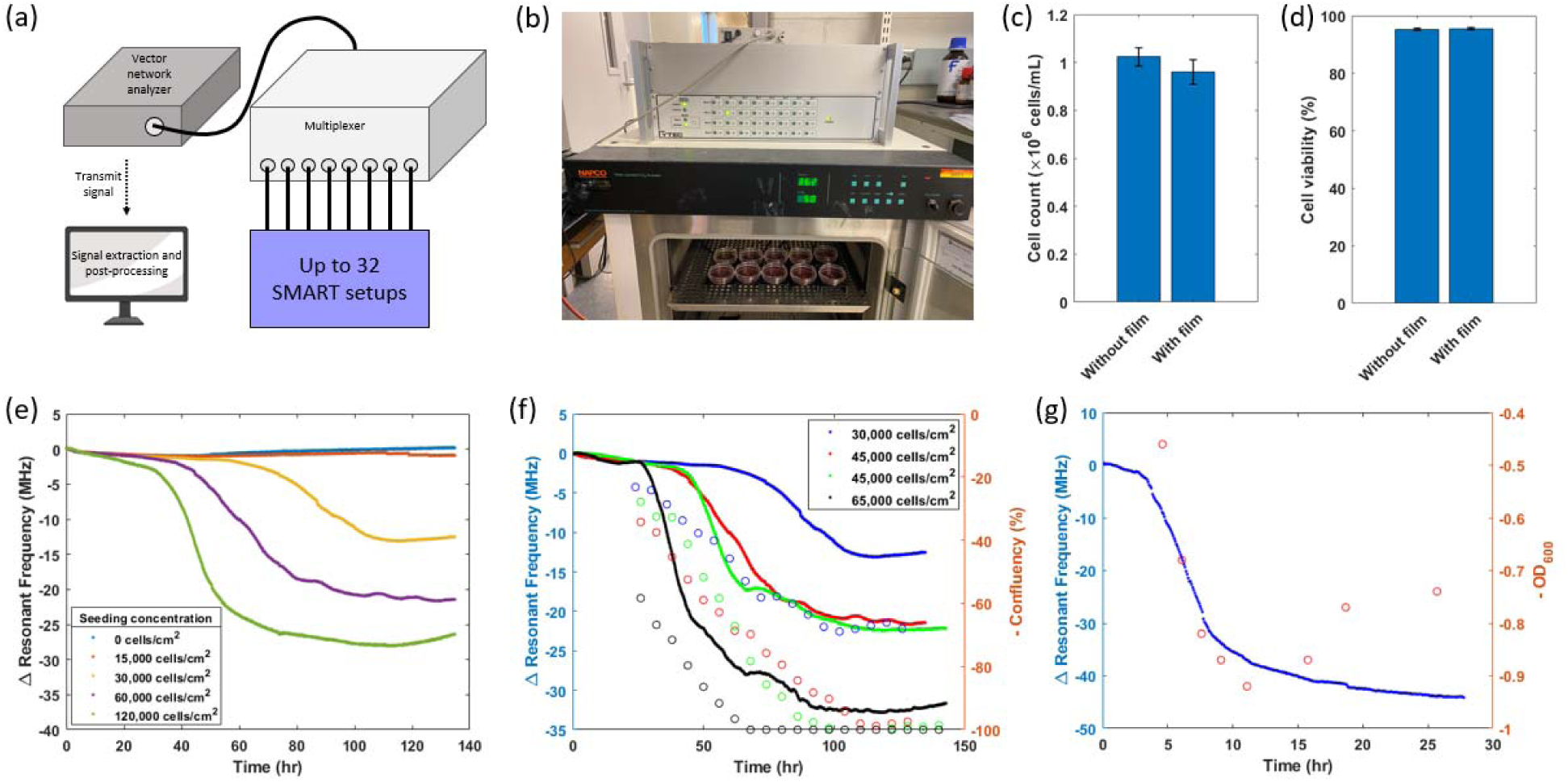
High throughput SMART setup and characterization of sensor response under different growth conditions and cell lines. (a) Schematic illustration of the high throughput, multiplexed SMART setup. (b) Image of operating SMART setup at high throughput. (c,d) Characterization of cell growth and viability of K562 cells in the presence of polyacrylate. (c) p-value = 0.161 (d) p-value = 0.722. (e) Resonant frequency profiles at different seeding densities. (f) Resonant frequency profiles at different seeding densities benchmarked to confluency. Lines indicate the continuous resonant frequency profile and circles show confluency from sampled images. Negative confluency values were used to match the resonant frequency trend. (g) Resonant frequency profiles for *E. coli* growth. Blue data points represent the continuous resonant frequency profile and red circles represent the *OD_600_* value. Negative of *OD_600_* values was used to match the resonant frequency trend.

To evaluate the SMART dish ability to transduce different growth rates, we varied the seeding concentration of HeLa cells (Figure 6e). We observed clearly distinguished response curves between the different seeding concentrations. At seeding concentration of 15 000 cells/cm^2^, the resonant frequency response did not vary greatly from the control with no cells. Microscopic images at the end of culture period showed little sign of growth, likely due to insufficient seeding concentration to support the growth. At seeding concentrations of 30 000, 60 000, and 120 000 cells/cm^2^, the resonant sensor generated 17, 21, and 27 MHz shifts, respectively, resulting in a gain of 0.105 MHz/1000 seeded cells/cm^2^ (Figure S11) More importantly, the resonant frequency profile at higher seeding concentration demonstrated a more rapid change in resonant frequency and plateaued more quickly. Similar results were also obtained with varying concentration of serum, in which higher serum concentration of serum resulted in a faster growth and thus the earlier rapid decrease in resonant frequency (Figure S12). These results suggest that the sensor prototype can differentiate not only the growth extent, but also the dynamic growth property between different cultures, a significant advantage over other, single time point, off-line proliferation assays.

To quantitatively compare the resonant frequency response to cell expansion, we calculated the confluency via manual annotation of HeLa cells microscopic images (Figure S13) during which the resonant sensor measurement was taken. As expected, both the resonant frequency and confluency profiles exhibited variable responses at seeding concentrations of 30 000, 45 000, and 60 000 cells/cm^2^ (Figure 6f). Note that the confluency profile was plotted in negative values to match the trend of resonant frequency profile. The confluency profiles showed a more linear and earlier increase than the fast change in resonant frequency, which indicates that the target metabolites are secreted during a more mature growth phase. In addition, the confluency and resonant frequency profiles plateaued almost consistently, especially the profiles for 30 000 cells/cm^2^. For the 30 000 cells/cm^2^ seeding concentration, the plateaued signal indicated nutrient depletion and this feature could be useful in bioprocess monitoring (signaling need for media change). At higher seeding concentrations, the confluency eventually achieved saturation and was unable to be further quantified using microscopic images. Since the automated segmentation of microscopic images has been used as a cell progression technique, the SMART system presented another modality with a higher detectable range and one that is not limited to monolayer cultures.

### Sensor response to other cell lines

Although the SMART setup demonstrated promising results using the HeLa cell line, it is critical to assess if the same metabolite secretion transduction method works with other common cell lines. Similar response curves were observed when tested with HEK293, K562, Jurkat, Chinese hamster ovary cells, and *Escherichia coli*. This indicates that the SMART dish could be applied to both eukaryotes and prokaryotes, as well as adherent and suspension cell types. We next attempted to benchmark the resonant frequency profile to the *E. coli* growth through *OD_600_* measurements. The resonant frequency entered the rapid change phase after about 4 hours of culture. Consistently, the *OD_600_* measurements showed that the rapid change in resonant frequency occurred during the exponential phase (Fig 6g). According to the *OD_600_* profile, the culture entered the stationary phase at around 11 hr. Although the resonant frequency profile was not able to distinguish the decrease in *OD_600_* due to the irreversible transduction mechanism, the resonant frequency profile plateaued around the stationary phase. Interestingly, when the same culture suspension (after resonant frequency plateaued) was transferred into a new SMART dish, the resonant frequency gradually decreased and eventually plateaued even though the cell culture did not further increase in *OD_600_*. As the bacteria are known for being resilient, it was likely that the bacteria were still dividing and secreting the metabolites despite no increase in *OD_600_* due to the rate of cell death compensating for the rate of cell growth.

### SMART signal consistency and characterization

After investigating the characteristics of the SMART system, we attempted to examine the sensor’s consistency and reliability by having multiple sensors (n=3) into a single 9 cm dish to remove the potential variability from different cultures and investigated the sensor consistency. We cultured K562 cells at seeding concentrations of 0.1, 0.167, and 0.5 million cells/mL. All three conditions resulted in the sigmoidal response as attained above (Figure 7a). The cell concentrations at the end of experiment were 1.1, 1.1, and 1.5 million cells/mL for 0.1, 0.167, and 0.5 million cells/mL seeding concentrations, resulting in a 11-, 6.6-, and 3-fold expansion, respectively. The maximum standard deviation (a measure of consistency) for 0.1, 0.167, and 0.5 million cells/mL seeding concentration was 5.3, 5.7, and 5.2 MHz, respectively. The highest standard deviation typically occurs around the exponential phase, possibly due to different diffusion and adhesive conformation patterns occurred between different sensors, as also noted by the better consistency before the exponential phase where less secretion occurs. Regardless, the SMART system can clearly distinguish between the different culture conditions. We next parameterized the SMART response curve by taking the first derivative (center finite difference) and fitting the rate with a Gaussian equation (Figure 7b). The five parameters are summarized in Table 1.

**Figure 7.**
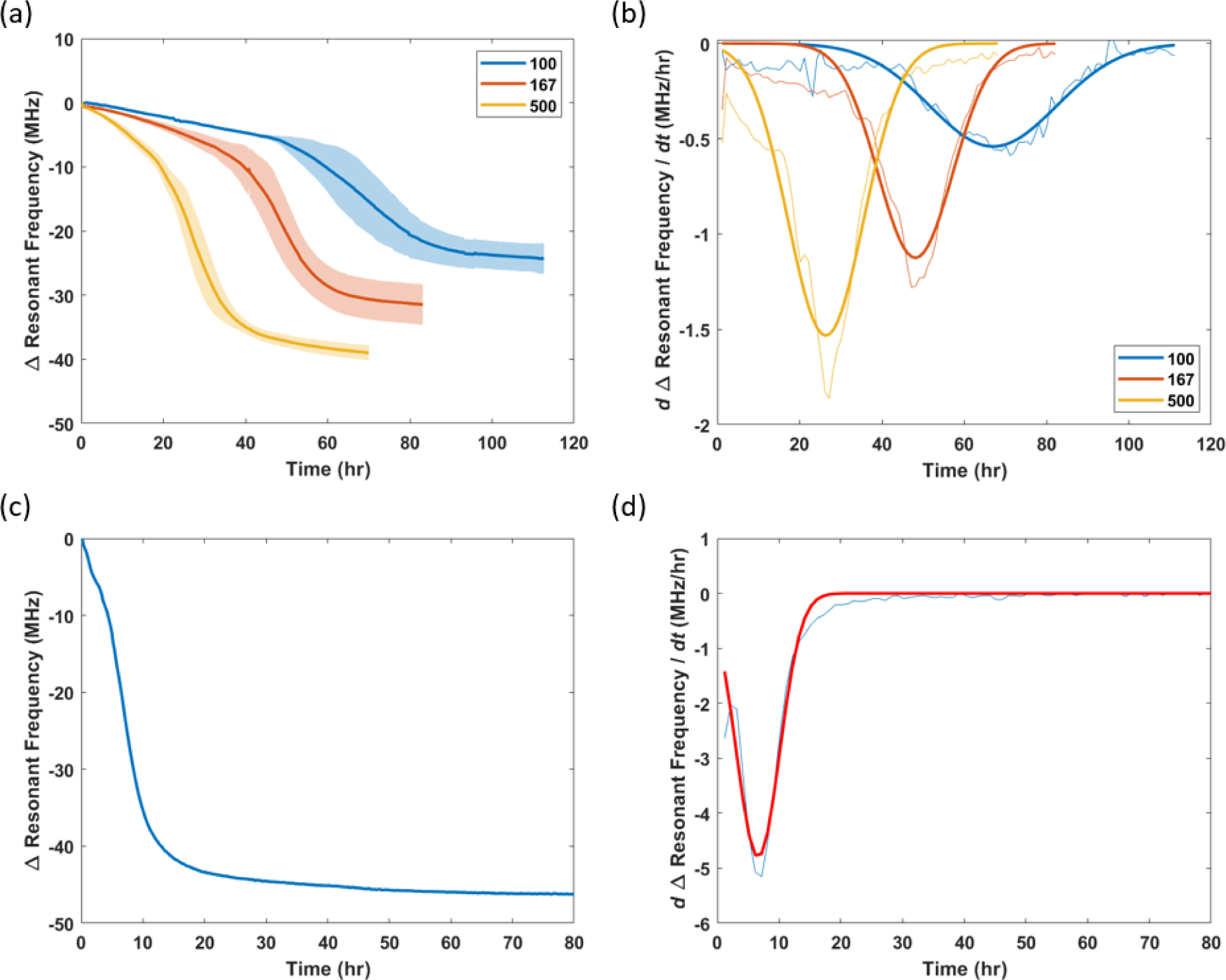
Reproducibility and reliability characterization and evaluation of SMART responses. (a) Resonant frequency profile at different K562 starting concentrations. (n=3). (b) First order derivative of resonant frequency profile from (a). Data was fitted with Gaussian function as shown in by the thicker lines. (c) Resonant frequency profile of a saturated, grown culture on a fresh SMART dish. (d) First order derivative of resonant frequency profile from (c). Data was fitted with Gaussian function as shown in by red line.

**Table 1.** Parameters from resonant frequency response curve.

| Parameters | Description |
| --- | --- |
| $\Delta f_f$ | Total change in resonant frequency. |
| $r_{max}$ | Maximum rate of change in resonant frequency.<br>Peak height of the Gaussian function fitted on the first derivative response curve. |
| $t_{max}$ | Time at maximum rate.<br>Peak position of the Gaussian function. |
| $t_{exp}$ | Duration of exponential change in resonant frequency.<br>Two times of the standard deviation of the Gaussian function. |
| $t_f$ | Time to reach resonant frequency plateau.<br>$t_f = t_{max} + t_{exp}$ |
| $A$ | Area of resonant frequency response curve. |

The parameters for each culture condition in Figure 6a,b was determined and tabulated in Table 2. The Δ*f_f_* increases with increasing K562 seeding concentration, resulting in a gain of 0.032MHz/1000 cells/mL (Figure S14) and indicating a higher metabolite synthesis that causes the culture media to alter the acrylic property at a higher degree. Note that the Δ*f_f_* is different between the 0.167 and 0.1 seeding concentrations but has a similar concentration at the end point, indicating that Δ*f_f_* is a result of cells secretion at the end point as well as the proliferation process. However, further investigation needs to be done due to the noise that could result from the SMART sensor and the hemacytometer. In addition, the different Δ*f_f_* suggests that the resonant frequency at equilibrium is determined by the metabolite concentration and not saturated by continuous accumulation. The *r_max_* increases with increasing seeding concentration, indicating that the secretion increases at higher cell concentration. The *t_max_* decreases with increasing seeding concentration, suggesting that the cultures enter the exponential phase in a shorter time. The *t_exp_* is longest at the lowest seeding concentration, probably due to a longer and slower proliferation process. Interestingly, *t_exp_* is similar between the 0.167 and 0.5 million cells/mL starting concentration. This parameter could therefore be used to indicate cell physiology where a longer exponential duration may indicate slower growth process and potentially unhealthy state (example shown in next section). The area of the response curve, *A*, decreases with increasing seeding concentration. This metric combines the resonant frequency shift, and cell growth time, *t_f_*. Although a higher seeding concentration resulted in a larger change in Δ*f_f_*, the short *t_f_* reduces the overall magnitude. One potential use case is in bioprocesses where the value is kept within an optimized range to maximize growth within the shortest time.

**Table 2.** Parameter analysis of K562 cell culture at different seeding concentrations.

| Cell seeding concentration<br>($\times 1000$ cells/mL) | Parameters | | | | | |
| --- | --- | --- | --- | --- | --- | --- |
| | $\Delta f_f$<br>(MHz) | $r_{max}$<br>(MHz/hr) | $t_{max}$<br>(hr) | $t_{exp}$<br>(hr) | $t_f$<br>(hr) | $A$<br>($\times 10^6$) |
| 100 | 24.3 | 0.54 | 67.0 | 43.6 | 110.6 | 1443 |
| 167 | 31.5 | 1.13 | 48.1 | 25.7 | 73.8 | 1337 |
| 500 | 39.0 | 1.53 | 26.4 | 25.9 | 52.3 | 1013 |

As we observed that Δ*f_f_* at equilibrium was not due to saturation of biomolecules within the polyacrylate film, we then investigated upper limit of *r_max_* by replacing the fresh media used to pre-equilibrate the sensor with a fully expanded, stationary phase K562 culture (Figure 6c,d). The *r_max_*, *t_max_*, and *t_exp_* were determined to be 4.8MHz/hr, 6.4 hr, and 10.2 hr, respectively. The *r_max_* is much lower than the effects from prenol (Figure 3d), suggesting that the diffusion process into the polyacrylate film is not limited by the high concentration of metabolites; instead, this indicates that Δ*f_f_* is indeed resulted from the accumulation of metabolites, providing information over the entire culture progression and justifying that spent media would not cause a significant response due to lack of metabolite secreting source.

### Drug screening application

To demonstrate the use of SMART dishes in drug discovery, the sensor response to marimastat during cell culture was studied. Marimastat is an inhibitor to a broad spectrum of matrix metalloproteinase as well as metalloprotease-disintegrins that have effects on cell proliferation by interrupting cell communications. The interrupted cellular function can result in reduced biological processes and was hypothesized to have a reduced metabolite secretion. At 20 hours post-inoculation, marimastat was administrated into four dishes at 0 (control), 7.5, 15, and 30 μM (final concentrations) and continued culturing for up to 4 days (Figure 8a). The marimastat resulted in 32.5, 25.2, 22.2, and 8.3 MHz Δ*f_f_* at 107 hours for marimastat concentrations of 7.5, 15, and 30 μM, respectively. Microscopic images at the end of culture validated that the introduction of marimastat resulted in reduced growth compared to the control. The results of the other parameters defined above are tabulated in Table 3. The *r_max_* decreased with increasing marimastat concentration. At 30 μM concentration, the resonant frequency did not result in a sigmoidal curve and microscopic images confirmed the minimal growth from this condition, suggesting that exponential growth was inhibited at this concentration. The *t_max_* increased with increasing marimastat concentration, indicating that marimastat delayed the exponential growth process due to slowing of normal growth. *t_exp_* first decreased at 7.5 μM and increased at 15 μM, indicating two ways of growth inhibition by shortening *t_exp_* or lengthening *t_exp_* but much lower *r_max_*.

**Figure 8.**
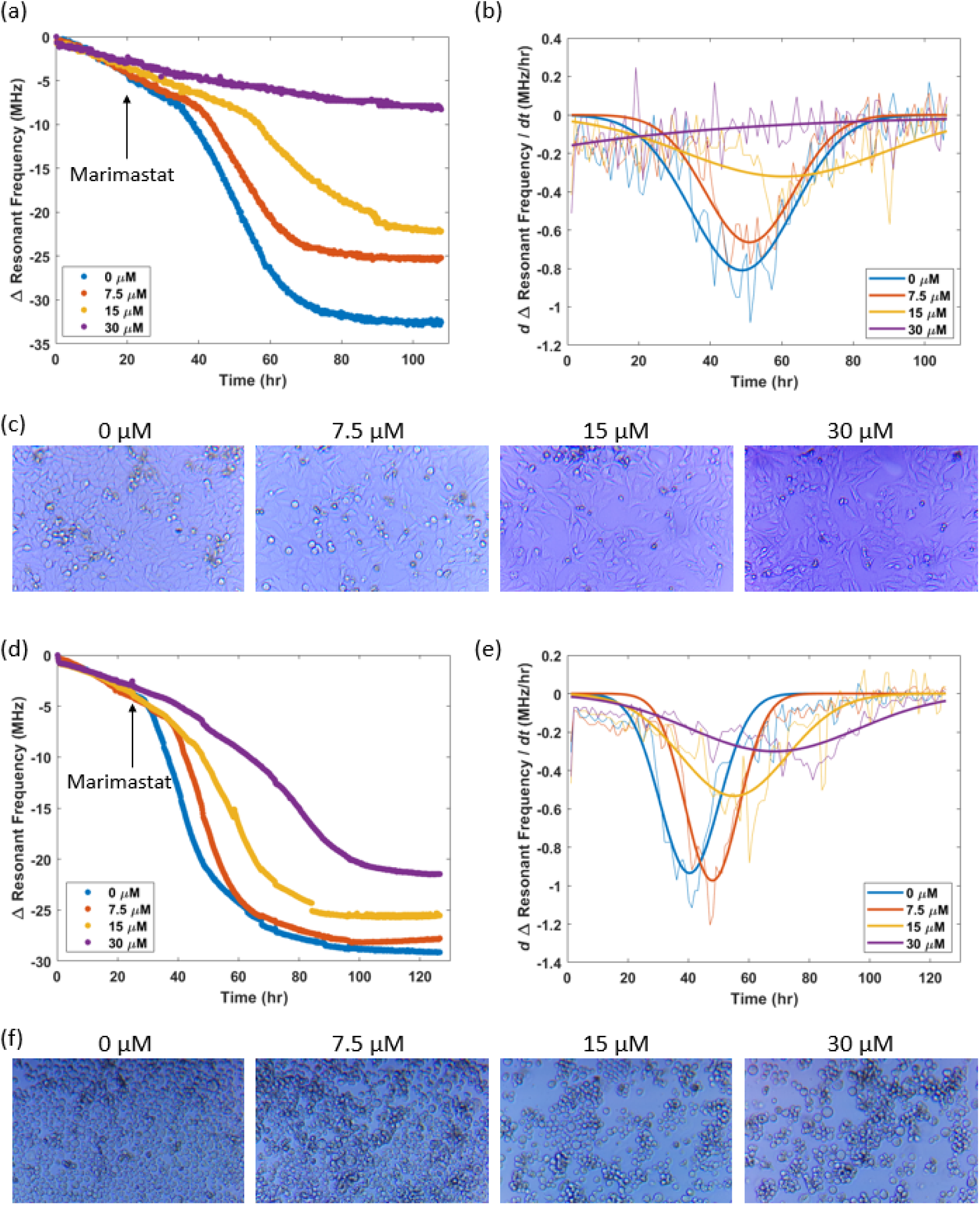
Drug screening using polyacrylate transduced resonant sensor. (a) Resonant frequency profiles of HeLa cells in response to marimastat concentrations of 0, 7.5, 15, and 30μM. (b) First derivative of resonant frequency profiles obtained in (a) fitted with Gaussian function. (c) Microscopic images of HeLa cells under different marimastat concentrations treatment after 107 hours of culture. (d) Resonant frequency profiles of K562 cells (seeding concentration = 0.2 ×10^6^ cells/mL) in response to marimastat concentrations of 0, 7.5, 15, and 30μM. (e) First derivative of resonant frequency profiles obtained in (d) fitted with Gaussian function. (f) Microscopic images of K562 cells under different marimastat concentrations treatment after 125 hours of culture.

**Table 3.** Parameter analysis of HeLa cell culture in treatment with marimastat.

| Marimastat concentration ($\mu\text{M}$) | Parameters | | | | | |
| --- | --- | --- | --- | --- | --- | --- |
| | $\Delta f_f$ (MHz) | $r_{max}$ (MHz/hr) | $t_{max}$ (hr) | $t_{exp}$ (hr) | $t_f$ (hr) | $A$ ($\times 10^6$) |
| 0 | 32.5 | 0.81 | 48.8 | 40.7 | 89.5 | 1484 |
| 7.5 | 25.2 | 0.66 | 51.0 | 34.9 | 85.9 | 1136 |
| 15 | 22.2 | 0.32 | 60.6 | 79.1 | 139.7 | 1192 |
| 30 | 8.3 | - | - | - | - | 331 |

A similar study was also performed using K562 cells at 0.2 million cells/mL seeding concentration. Marimastat was introduced at 24h post-inoculation and the resulting response curve parameters are shown in Table 4. In general, similar effects with the treatment on HeLa cells were obtained. The cell counts and viability at the end for treatment concentrations of 0, 7.5, 15, and 30 μM were 1.41, 1.42, 0.97, 0.40 million cells/mL and 92, 92, 90, and 91 %, respectively. High cell viability indicates that the resulting response profiles metalloprotease inhibition reduces the cell proliferation but less effect on the metabolite secretion.

**Table 4.**
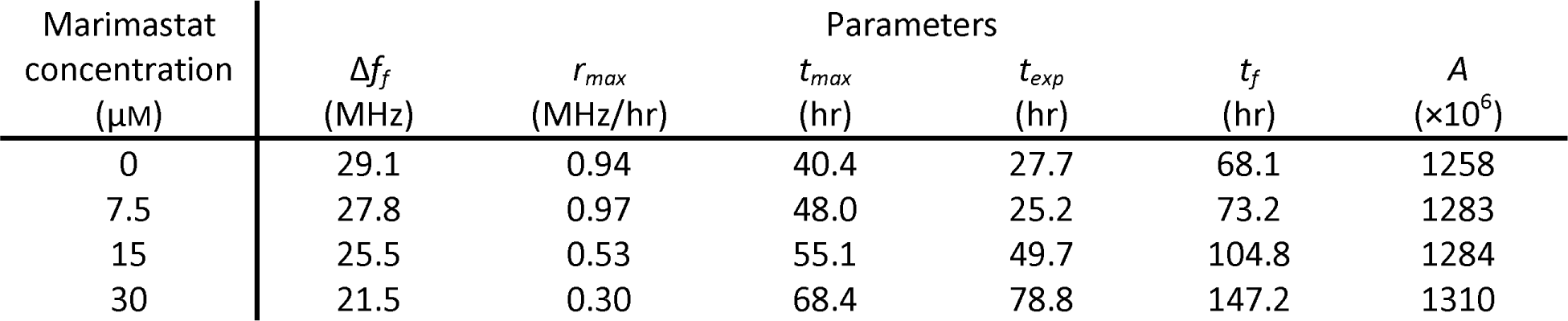
Parameter analysis of K562 cell culture in treatment with marimastat.

### SMART Dish future directions

The SMART system demonstrates a unique capability of in-line monitoring of cell culture progression. Although we performed experiments mostly using culture dishes, the sensor can also be applied to other culture flasks including static breathable vessels (e.g. G-Rex), well-plates, T-flasks, shaker flasks, etc. However, the SMART system may encounter difficulty in culture vessels with thicker walls or made with metal due to the inductive coupling limitation between the sensor and reader antenna. Additionally, the current SMART setup requires up to 12 hours of pre-incubation with culture media before cell addition to achieve equilibrium and a stable, baseline resonant frequency. Although this resonant frequency shift in the beginning has been observed, the shift has been consistent and can be eliminated by background subtraction. Biological pathways responsible for the plasticizing metabolites remain to be thoroughly investigated, which can then be used for analysis of biological processes within a live cell. Chemical modification of polyacrylate or other polymer materials to adjust the affinity towards other biomarkers opens the possibility to investigate other biological parameters.

## Conclusion

A new modality of continuous metabolite monitoring via affinity-based absorption of polymers has been discovered. Secreted metabolites can be used to track biological processes such as cell growth; however, current metabolite monitoring strategies rely on intermittent sampling and are not well-suited for continuous monitoring. Herein, we utilized the affinity of cross-linked polyacrylate to molecules containing C=C and O-H functional groups like secreted terpenoids; these plasticize the polymer in a concentration-dependent manner. By coupling this membrane with resonant sensors to transduce the mechanical changes to shifts in resonant frequency, we demonstrated SMART dishes that enable continuous monitoring of metabolites within closed vessels through non-destructive, continuous metabolite absorption. The SMART system resulted in over a 30 MHz resonant frequency shift, over 20-fold increase in sensitivity compared to sensing through the intrinsic cell permittivity. The SMART dish signal was successfully correlated to cell proliferation and interrupted cell growth. The dynamic response curve subsequently enables the extraction of multiple parameters that indicate culture conditions such as starting cell concentration and inhibitory drug effects. The wireless interface, scalability, and high acquisition rate should enable applications in pharmacological discovery as well as biomanufacturing.

## Materials and Methods

### Materials

HeLa cells were obtained from Prof. Surya Mallapragada’s laboratory (Iowa State University). Jurkat and K562 cells were obtained from ATCC. Liposomes were obtained from Prof. Rizia Bardhan’s laboratory (Iowa State University).

Polyacrylate adhesive (3M 467MP) was obtained from Grainger. Phenol, menthol, geraniol, trigonelline hydrochloride, pyridine hydrochloride, quinine hemisulfate salt monohydrate, erythromycin, and 1-hydroxyanthra-9,10-quinone were purchased from Sigma.

### SMART fabrication

Resonant sensors were fabricated either using chemical etching or wire wound methods. Fabrication through chemical etching method was described in previous work. Briefly, a mask was first created on a copper laminate (Pyralux) using a solid ink printer or a lithography method. The unmasked copper was removed in an etching solution containing a 1:1 volume ratio of 3 wt% hydrogen peroxide and concentrated hydrochloric acid. Finally, the mask was removed by acetone rinsing. For SMART setups, wire wound resonant sensor was used to enable sufficient air voids. The wire wound resonant sensor was fabricated using an enameled copper wire of 18 cm, 32 AWG and a spiral making tool. The fabricated resonant sensor was then immobilized onto a polyacrylate adhesive in a culture dish. Another polyacrylate adhesive was applied onto the resonant sensor, creating a polyacrylate-resonant sensor-polyacrylate layered structure. The SMART dishes were sterilized using 70% ethanol and air-dried in a UV chamber before use.

### High-throughput resonant sensor readout system and microscope imaging system

A vector network analyzer (Copper Mountain Technologies TR1300) was used to measure the reflectance over a frequency range. For high-throughput measurements, a multiplexer (Cytec) was used to connect multiple readout coils and the vector network analyzer. The readout coil was fabricated using a single loop, 20 AWG copper wire of the same diameter as the resonant sensor. A holder for the readout coil was fabricated by 3D printing (Ultimaker S3) using acrylonitrile butadiene styrene (ABS) filament. For microscopic image acquisition, a path is created in the center of the readout coil holder to allow light penetration through the magnification lens. A 20× magnification lens in combination with a camera (Raspberry Pi High Quality Camera, 12.3 megapixel) was programmed to capture images at a 15 minute interval. To obtain confluency, the pixels containing cells were manually annotated. The confluency was determined by the number of pixels containing cells divided by the total number of pixels. For the reflectance measurement, the spectrum in the frequency range of 10 MHz to 200 MHz containing 5000 points was obtained every 5 minutes. Background subtraction was performed prior to the resonant frequency extraction.

### Cell culture

HeLa cells were cultivated in DMEM supplemented with 10% (v/v) FBS and 1% (v/v) penicillin-streptomycin. Jurkat, CHO, and K562 cells were cultivated in RPMI-1640 supplemented with 10% (v/v) FBS and 1% (v/v) penicillin-streptomycin. E. coli BL21 cells were cultivated in M9 media (Sigma) supplemented with 2% (v/v) of 1M glucose and 0.2% (v/v) of 1M magnesium sulfate. All cells were cultured in a humidified atmosphere at 37°C and 5% CO_2_, except e. coli was cultured without CO_2_. Media exchange (media addition for suspension cells) was performed every other day, but not during the experiment unless specifically stated.

Cell counting was mainly performed using hemocytometer except for the study of the polyacrylate effect on cell growth (Figure 6c). Cell suspension was mixed with trypan blue in 1:1 volume ratio for viability assessment. For the study of the polyacrylate effect on cells, K562 cells were seeded at 0.3 million cells/mL into well plates containing polyacrylate films (n=3) and without polyacrylate films (n=3) and cultured for 4 days. After 4 days, counting beads (Invitrogen™ C36995) was added into the cell suspension to a final concentration of 0.48 ×10^5^ beads/mL and counted using a flow cytometer (FACSCanto).

### Chemical screening

Organic chemicals with poor solubility were dissolved to saturation in ethanol (see Table S1 for details). Prior to the addition of each chemical, the SMART sensor was pre-equilibrated in water or culture media for about 20 hours or until the signal stabilized under 37°C, humidified environment.

### Polyacrylate topography validation

A polyethylene terephthalate (PET) of 0.1 mm thickness was laser cut (Glowforge) at a speed setting of 300 and at full power, creating wells of about 0.2×10 mm^2^. An adhesive transfer tape was marked with a straight line and applied onto the PET film (with the marked side of the adhesive in contact with the film). The PET matrix was transferred to a culture dish, sterilized with ethanol, and proceeded with culturing of HeLa cells or remained culture media (control) for 4 days. After 4 days, the cells were removed with trypsin and the height profile of the matrix was analyzed using a digital microscope (Olympus DSX 110).

### Characterization

The polyacrylate adhesive was cut into 2.5×2.5 cm^2^ slices and placed into a petri dish containing only media or expanded K562 culture (1.5×10^6^ cells/mL) for at least 2 days. The polyacrylate film was gently placed on the media surface while the cells were suspended at the bottom to reduce the likelihood of cells embedding into the polyacrylate. After 3 days, the polyacrylate films were rinsed with deionized water and analyzed without drying.

The polyacrylate-containing adhesive was characterized by FTIR spectroscopy to track evidence of chemical changes upon exposure to cell metabolites using an Attenuated Total Reflection (ATR) FTIR spectrometer equipped with an (iD7-ATR accessory, Thermo Fisher Scientific Nicolet^TM^iS^TM^ 5 Spectrometer). The spectral range of observation was 4000-400 cm^-1^ with a resolution of 8cm^-1^, collecting 64 scans per sample.

Rheological properties of polyacrylate adhesive were characterized using a rheometer (ARES-G2, TA Instrument). Two slices of treated polyacrylate films were compressed and analyzed using an 8 mm parallel plate. The frequency was set at 1 Hz and the temperature was set at 35°C.

Thermal characterization of the polyacrylate adhesives was carried out by differential scanning calorimetry performed on the DSC-Discovery 2500 series (TA instruments). 4-7 mg of the polyacrylate samples were weighed and hermetically sealed in aluminum pans. The method involved heating the samples from -80C to 250C at a ramp rate of 10C/min under constant nitrogen sparge maintained at 50ml/min.

Thermal stability was characterized by TGA (STA 449 F1 NETZSCH).

For characterization of embedded molecules, the treated polyacrylate films after the preparation steps were mixed with 6 mL of acetone for 1 hr at 37°C. After swelling, 500 μL of the acetone solution was extracted and analyzed using LCMS (Agilent 6540 LC-Q-TOF).

### Statistical analysis

Data with replicates are shown as the mean values ± standard deviation. Error bars represent the standard deviation. Significance or p-value was computed using unpaired t-test.

## Acknowledgements

We acknowledge helpful discussions and feedback from Professor Ian Schneider, Professor Marit Nilsen-Hamilton, Professor Surya Mallapragada, and Professor Metin Uz during the course of this work.

