## Supplementary material for "Single-use, Metabolite Absorbing, Resonant Transducer (SMART) culture vessels for label-free, continuous cell culture progression monitoring": Figure S

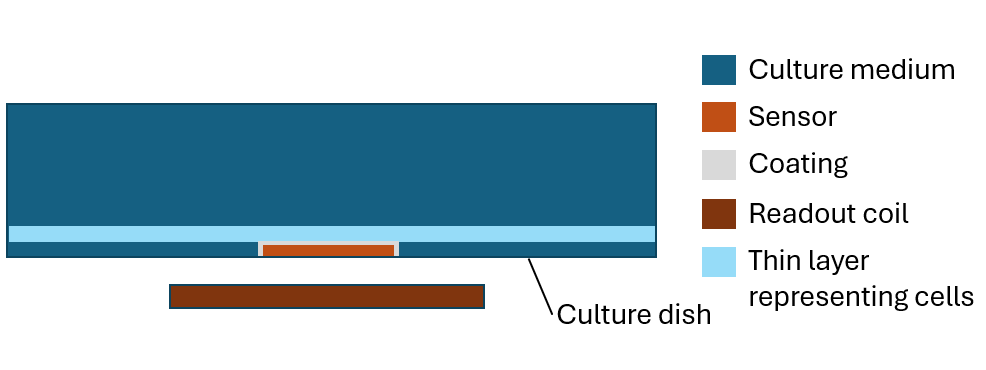

Figure S1. Cross sectional side view of simulation of the resonant sensor system to investigate sensitivity to changing permittivity of a thin layer (not to scale). Thin layer height: 3μm; Dish height: 10mm; Sensor height: 10μm.

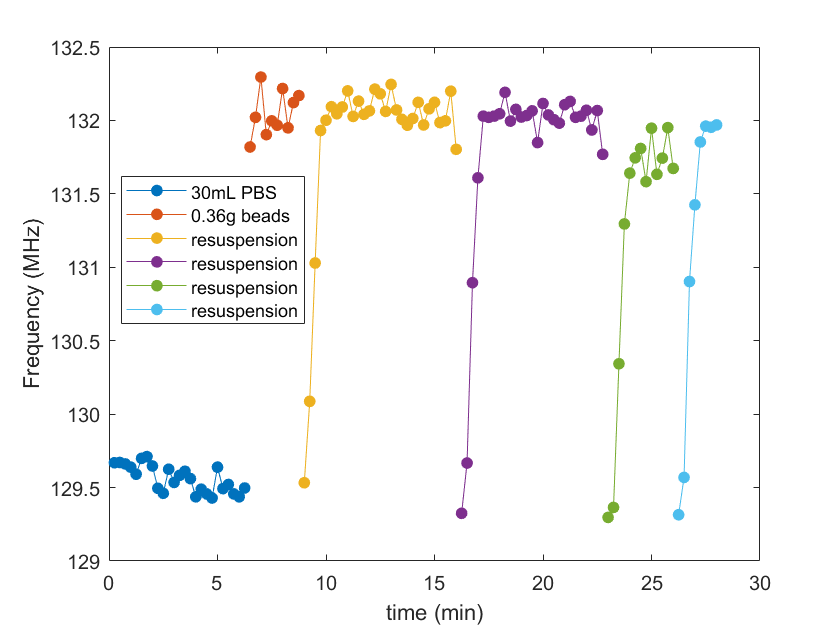

Figure S2. Sensitivity of a wire wound resonant sensor of length 23 cm to polyethylene beads during resuspension events. Resonant sensor was insulated with a 63.5μm Kapton.

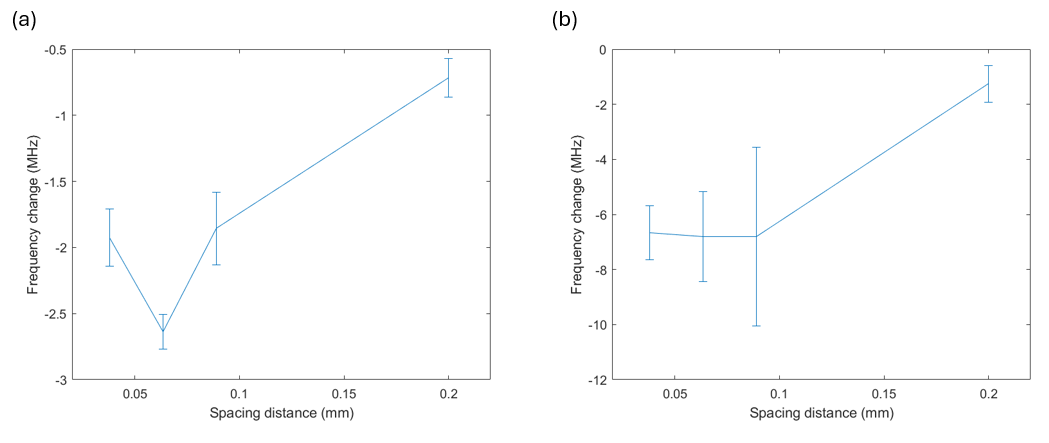

Figure S3. Sensitivity of resonant sensor to suspending polyethylene beads with (a) 23 cm wound wire and (b) 15 cm wound wire.

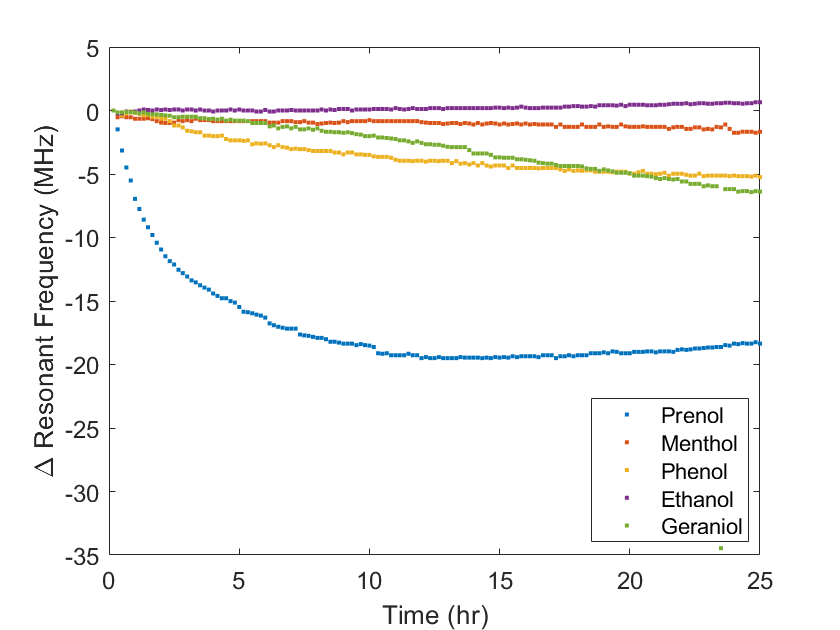

Figure S4. Screening of molecules containing different functional groups.

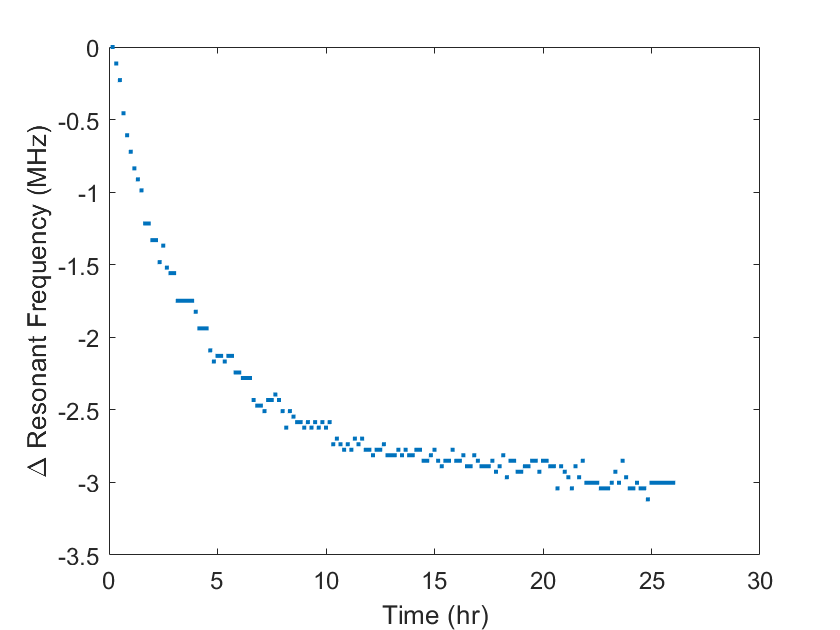

Figure S5. SMART equilibrium response to the initial addition of culture media.

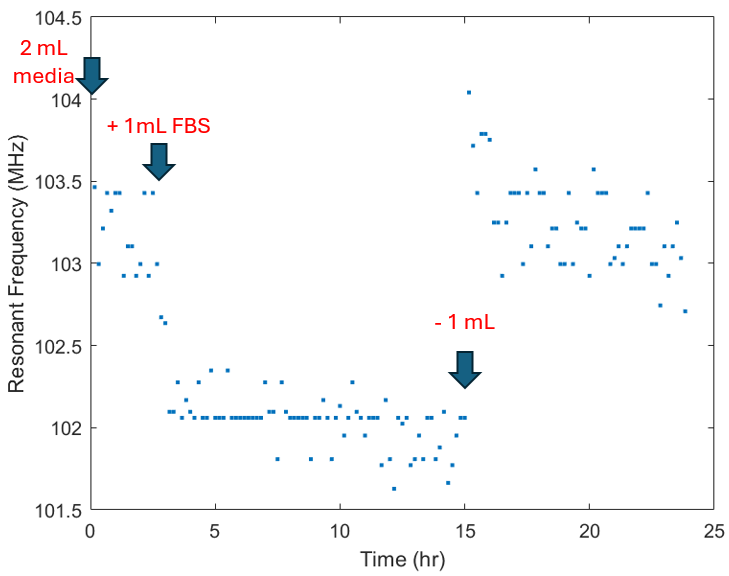

Figure S6. SMART response to fetal bovine serum (FBS). SMART was pre-equilibrated with culture media. Signal response after FBS addition is mainly due to increased media volume, as noted by return of signal upon solution removal.

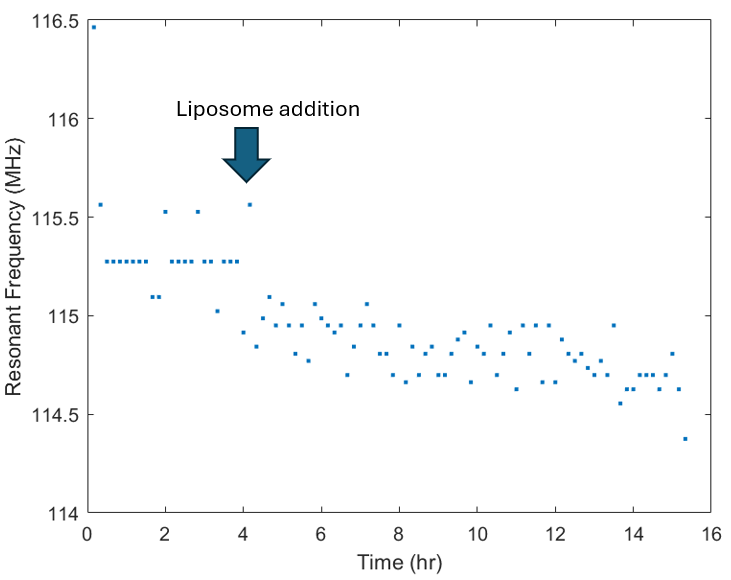

Figure S7. SMART response to liposome. SMART was pre-equilibrated with culture media.

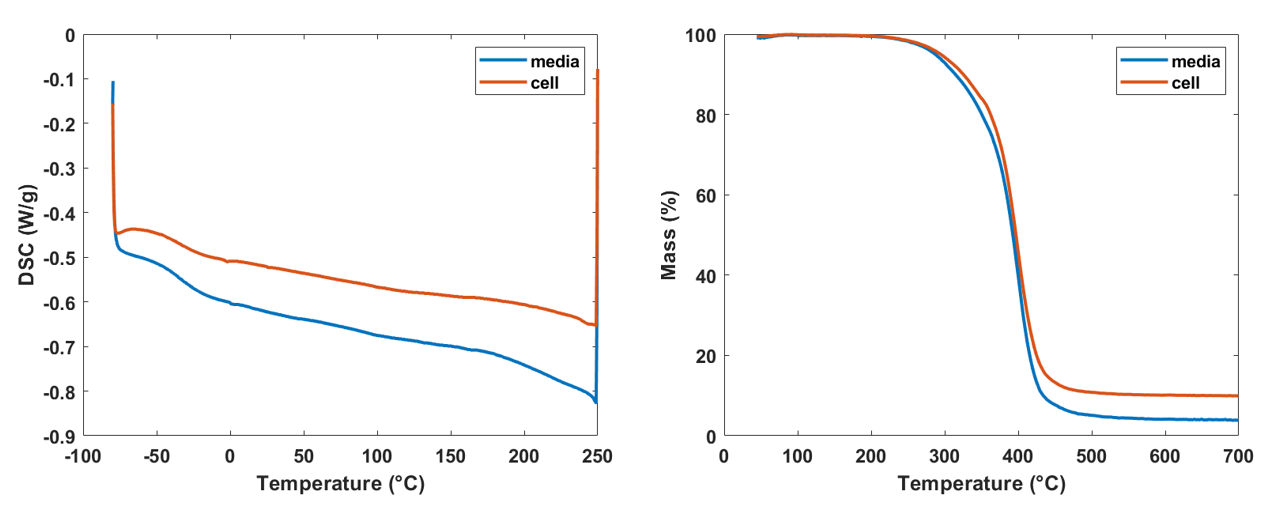

Figure S8. (Left) Differential Scanning Calorimetry and (Right) Thermogravimetric Analysis of polyacrylate after treatment with media or cells suspension.

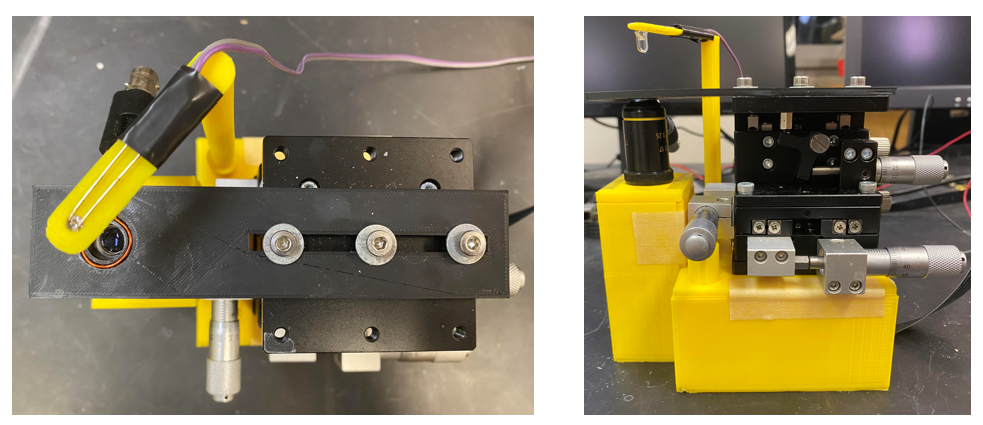

Figure S9. (Left) Top view and (Right) Side view of the frugal microscopic imaging system integrated with resonant sensor readout antenna.

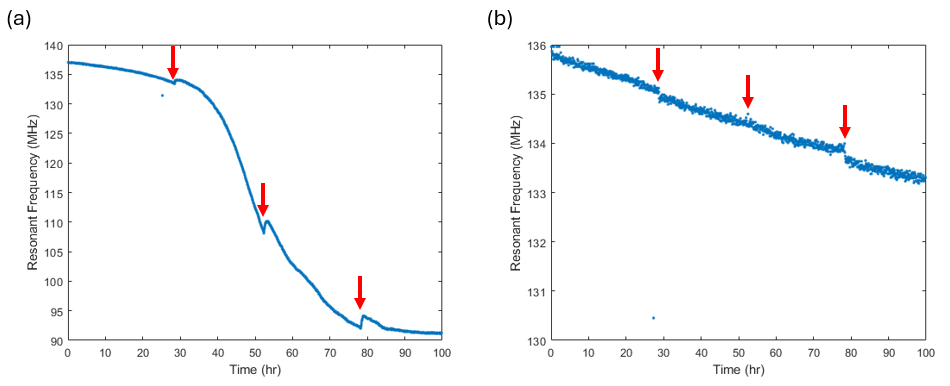

Figure S10. (a) Resonant frequency profile with media exchange at time points indicated by red arrows. (b) Resonant frequency profile of SMART dish containing only fresh media or spent media from (a) without cells.

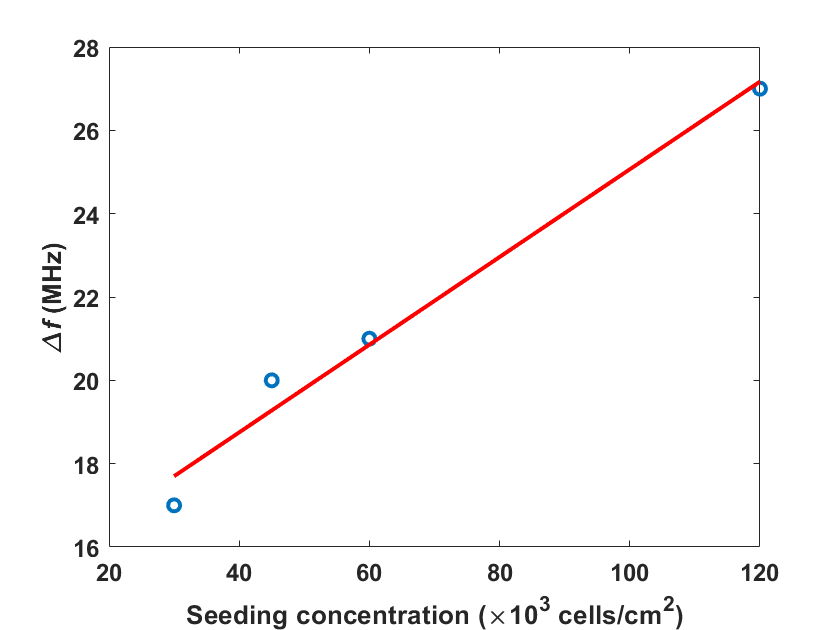

Figure S11. Correlation between resonant frequency change and seeding concentration of HeLa cells. Gain = 0.105MHz/1000cells/cm^2^. R^2^ = 0.98.

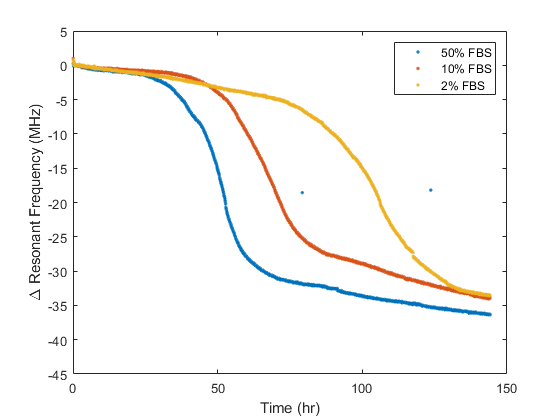

Figure S12. Resonant frequency profiles of HeLa cell culture under different concentrations of fetal bovine serum.

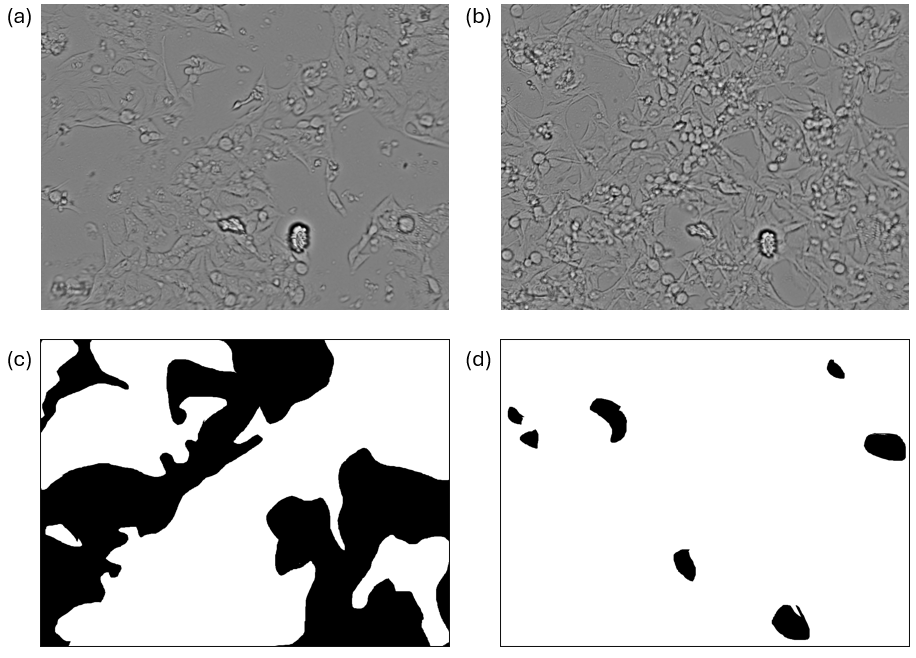

Figure S13. (a), (b) Microscopic images of HeLa cells at different time of culture and (c), (d) corresponding annotated images. White pixels represent cells whereas black pixels represent background.

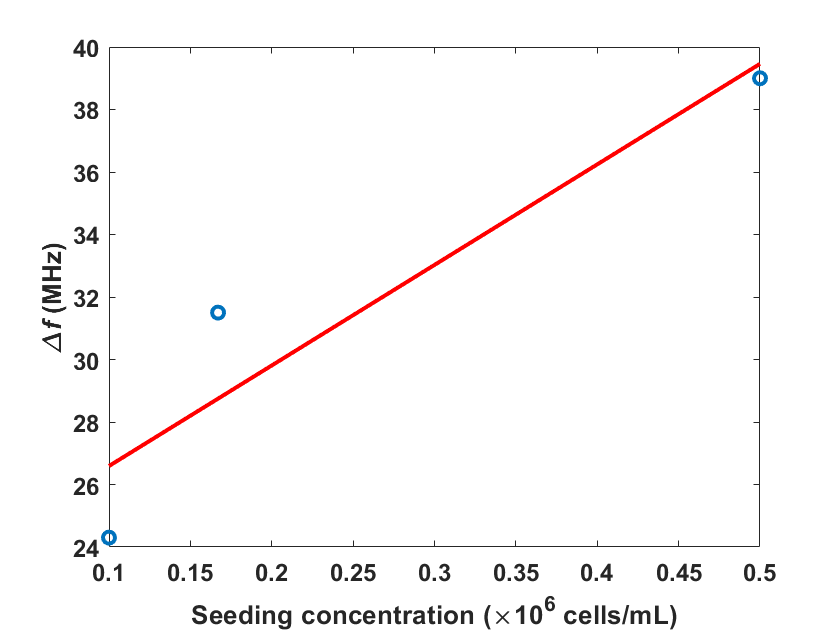

Figure S14. Correlation between resonant frequency change and seeding concentration of K562 cells. Gain = 0.032MHz/1000cells/mL. R^2^ = 0.88.

### Table S1. Formulation of screening solutions

| **Chemical** | **Solvent** | **Concentration** | **Volume added (7mL total volume)** |
| --- | --- | --- | --- |
| Lactic acid | - | >90 wt% | 1 μL |
| Ethanol | - | 100 % | 700 μL |
| Phenol | Ethanol | 200 mg/mL | 700 μL |
| Quinine hemisulfate | Ethanol | 8.6 mg/mL | 700 μL |
| Pyridine hydrochloride | Ethanol | 50 mg/mL | 700 μL |
| Trigonelline hydrochloride | Water | 50 mg/mL | 700 μL |
| Hydroxyanthraquinone | Ethanol | 2 mg/mL | 700 μL |
| Erythromycin | Ethanol | 50 mg/mL | 700 μL |
| Amphotericin | Water | 250 μg/mL | 700 μL |
| Menthol | Ethanol | 100 mg/mL | 700 μL |
| Geraniol | - | 98% | 700 μL |
| Prenol | - | 99% | 700 μL |
| Spent media | - | - | Replaced with 7 mL |
| Fetal bovine serum | - | - | 1 mL |
| Liposome | Phosphate-buffered saline | 4:1 Dipalmitoylphosphatidylcholine/  Cholesterol | 500 μL (2.5 mL total volume) |
